# COP9 signalosome complex subunit-7-mediated regulation of cAMP levels contributes to autophagic degradation and pathogenesis of rice blast fungus *Magnaporthe oryzae*

**DOI:** 10.1101/2023.04.25.538259

**Authors:** Lili Lin, Hengyuan Guo, Wajjiha Batool, Lianyu Lin, Jiayin Cao, Qiuli An, Sami Rukaiya Aliyu, Jiandong Bao, Zonghua Wang, Justice Norvienyeku

**Author notes:** Corresponding authors: Norvienyeku Justice and Zonghua Wang College of Plant Protection, Hainan University, Haikou 570228, China.

## Abstract

- Photo-dependent processes, including circadian rhythm, autophagy, ubiquitination, neddylation/deneddylation, and metabolite biosynthesis, profoundly influence microbial pathogenesis. Although a photomorphogenesis signalosome (COP9/CSN) has been identified, the mechanism by which this large complex contributes to pathophysiological processes in filamentous fungi remains unclear.
- Here, we identified eight CSN complex subunits in the rice blast fungus *Magnaporthe oryzae* and functionally characterized the translocon subunits containing a nuclear export or localization signal (NES/NLS).
- Targeted gene replacement of these CSN subunits, including *MoCSN3*, *MoCSN5*, *MoCSN6*, *MoCSN7*, and *MoCSN12*, attenuated vegetative growth and conidiation in *M. oryzae* and rendered non-pathogenic deletion strains. *MoCSN7* deletion significantly suppressed arachidonic acid catabolism, compromised cell wall integrity, subverted photo-dependent ubiquitination, and abolished photo-responsiveness. Surprisingly, we also discovered that MoCSN subunits, particularly MoCsn7, are required for the cAMP-dependent regulation of autophagic flux.
- Therefore, MoCSN significantly contributes to morphological, physiological, and pathogenic differentiation in *M. oryzae* by fostering cross-talk between multiple pathways.

## Introduction

Constitutive photomorphogenesis signalosome 9 (COP9 /CSN) is a multi-protein complex that profoundly influences diverse developmental processes in almost all living organisms(Barth et al., 2016). COP9/CSN modulates cellular, metabolic, and signal transduction pathways(Füzesi-Levi et al., 2020; Shackleford and Claret, 2010). The COP9 holocomplex in eukaryotes typically consists of 8-9 subunits. However, the number of subunits constituting a complex may vary among organisms. For instance, while the CSN complex in *Schizosaccharomyces pombe* consists of only four subunits, including CSN1 (Caa1 and Sgnsp), CSN2, and CSN4, the CSN complex in *Drosophila melanogaster*, *Arabidopsis thaliana*, and *Homo sapiens* is made up of seven, eight, and ten subunits, respectively(Gutierrez et al., 2020; Peng et al., 2001). The CSN complex was first identified as a vital photomorphogenesis machinery downstream of photoreceptors and has been shown to promote light-dependent physiological development in *A. thaliana* by modulating the nucleocytoplasmic distribution of COP1 in response to photoperiods(Osterlund et al., 1999). In addition to acting as negative regulators of photomorphogenesis in plants, subunits of the CSN complex have been implicated in several developmental and cellular processes, including ubiquitination, cell cycle regulation, metabolism, stress tolerance, reproduction, signal transduction, transcriptional regulation, and survival across different organisms(Shackleford and Claret, 2010).

The CSN signalosome complex supports numerous developmental events in eukaryotes, partly by facilitating ubiquitin-dependent degradation of aged, misfolded, and mislocalized proteins and promoting ubiquitin homeostasis(Kim et al., 2022). For example, studies have shown that selected subunits of the CSN complex directly interact with the 19S regulatory unit of the 26S proteasome complex in plants and animals(Yu et al., 2011). The binding of these selected CSN subunits, including (CSN2/CSN6) and (CSN4/CSN5) to Cullin-RING E3 ubiquitin ligases of the SkpA-Cul1-F-box (SCF) complex in animals and plants results in deactivation of Cullin-RING E3 ubiquitin ligases by inhibiting or displacing Nedd8 from Cullin. The displacement of Nedd8 or deneddylation triggered by CSN-SCF interaction results in the termination of Cullin-dependent ubiquitination. Deneddylation is an essential biochemical process that effectively prevents auto-ubiquitination by coordinating interactions between substrates and substrate adaptor subunits of the SCF complex(Suisse et al., 2018).

The transcriptional repressor Capicua (CIC) promotes the proper development of organs (embryo, lungs, abdominal wall, and brain), guides cell differentiation (neural stem cells and T cells), and enhances the enterohepatic circulation of bile acids in mammals(Lee, 2020). Activation of the Epidermal Growth Factor Receptor and its downstream elements is essential for CIC mislocalization, Cullin 1-dependent ubiquitination, and timely degradation following CIC phosphorylation by MAP kinase(Chen et al., 2021). Recent studies have demonstrated that CSN-mediated deneddylation events impair Cullin1-dependent ubiquitination and CIC degradation, resulting in tumorigenesis and the development of various cancers. Individual CSN complex subunits regulate various cellular and developmental processes in eukaryotes(Serino and Deng, 2003). In Drosophila, disruption of *CSN4 and CSN*5 triggers a DNA double-strand-break-dependent meiotic checkpoint, resulting in larval mortality, attenuation of oocyte development, and photoreceptor differentiation(Cope and Deshaies, 2003). However, the deletion of *S. pombe CSN1* and *CSN2* arrested the progression of the cell cycle at the G2 phase, triggered the accumulation of damaged DNA, and rendered the *CSN1* and *CSN2* defective strains vulnerable to radiation-induced stress.

RNAi-mediated silencing of selected subunits of the CSN complex in *Caenorhabditis elegans* suppressed the activities of germ-line RNA helicases (GLHs) and katanin (a microtubule-severing factor), causing worm sterility and disrupting the functionality of multiple cellular processes, including microtubule polymerization, nuclear positioning, progression of cell division (anaphase), and cytokinesis(Hartman et al., 1998; Moghe et al., 2011; Pintard et al., 2003; Smith et al., 2002).

Compared with animal and plant orthologs, little is known about how filamentous fungus CSN complex subunits contribute to pathophysiological development. However, the impact of CSN-dependent regulation on photo-responsive processes, including circadian rhythm, morphogenesis, and sporogenesis, has been documented in fungi(Busch et al., 2003; Wilson et al., 2021). The photomorphogenesis signalosome complex helps maintain the periodicity of the circadian clock by facilitating the ubiquitination of the WD-40 domain-containing protein (FWD-1), a substrate-recruiting subunit of the SCF complex that activates the degradation of the essential circadian protein FREQUENCY (FRQ) following phosphorylation(He et al., 2005). Additional studies have shown that deletion of subunit-2 (CSN2) of the CSN complex in *Neurospora crassa* suppresses FRQ protein degradation and subsequently triggers indeterminate conidiation and attenuation in the vegetative growth of *N. crassa*(He et al., 2005). In *Aspergillus nidulans*, targeted gene deletion of CSN signalosome complex subunits disrupted the progression of photo-responsive developmental processes, deactivated regulatory mechanisms associated with secondary metabolic pathways, and abolished photosporogenesis in *A. nidulans* specifically lacking *CSN4* and *CSN5*(Busch *et al*., 2003). Furthermore, the deletion of *CSN5*, which is thought to catalyze deneddylation, in the necrotrophic plant pathogenic fungus *Alternaria alternata* impaired a series of developmental and cellular processes, including vegetative growth, conidiogenesis, stress tolerance, and virulence, and significantly downregulated the expression of genes associated with essential metabolic pathways(Wang et al., 2018).

Although CSN-regulated parameters such as sporulation, secondary metabolism, stress tolerance, ubiquitination, morphogenesis, cell division, and cell proliferation profoundly influence the dissemination, initiation, and development of diseases caused by filamentous plant pathogenic fungi, there are limited reports on the direct contributions of individual subunits of the CSN complex to the physiological and pathogenic development of phytopathogenic fungi. A limited understanding of the significance of CSN subunits in fungal pathogenesis is a bottleneck hindering the targeting of CSN signalosome complex subunits in antifungal compound development. The aim of this study was to identify core subunits that potentially regulate the nucleocytoplasmic shuttling of the CSN complex in *Magnaporthe oryzae*, a model fungus for studying plant-pathogen interactions, using bioinformatics analyses followed by targeted gene replacement, proteomics, and metabolomics. Such approaches will help to comprehensively evaluate the influence of CSN complex-mediated photomorphogenesis on photogenic processes, including sporulation, redox homeostasis, autophagy, and pathogenicity in the economically destructive rice blast fungus.

## Results

### *M. oryzae* CSN signalosome complex subunits share a close evolutionary relationship with *Fusarium graminearum* and *N. crassa* orthologs

Comparative phylogenetic analyses were performed to ascertain the evolutionary link between the individual CSN subunits identified in rice blast fungus and orthologs from *A. thaliana, A. nidulans, Aspergillus oryzae, F. graminearum, N. crassa, H. sapiens, O. sativa (ssp. japonica)*, and *Phytophthora infestan.* The results showed that in addition to Csn12, which is selectively conserved in *A. oryzae*, *A. nidulans, F. graminearum, M. oryzae*, and *N. crassa*, the canonical CSN subunits were conserved in all nine organisms sampled for phylogenetic studies. We noted that the evolutionary lineage of the individual CSN subunits identified in *M. oryzae* varies among organisms. For instance, while Csn1 and Csn5 in *M. oryzae* share evolutionary ties with orthologs in *N. crassa*, Csn4 aligns closely with its ortholog from *A. nidulans.* Csn6, Csn7, and Csn12 clustered with *N. crassa* and *F. graminearum* (Figure 1A-H).

**Figure 1.**
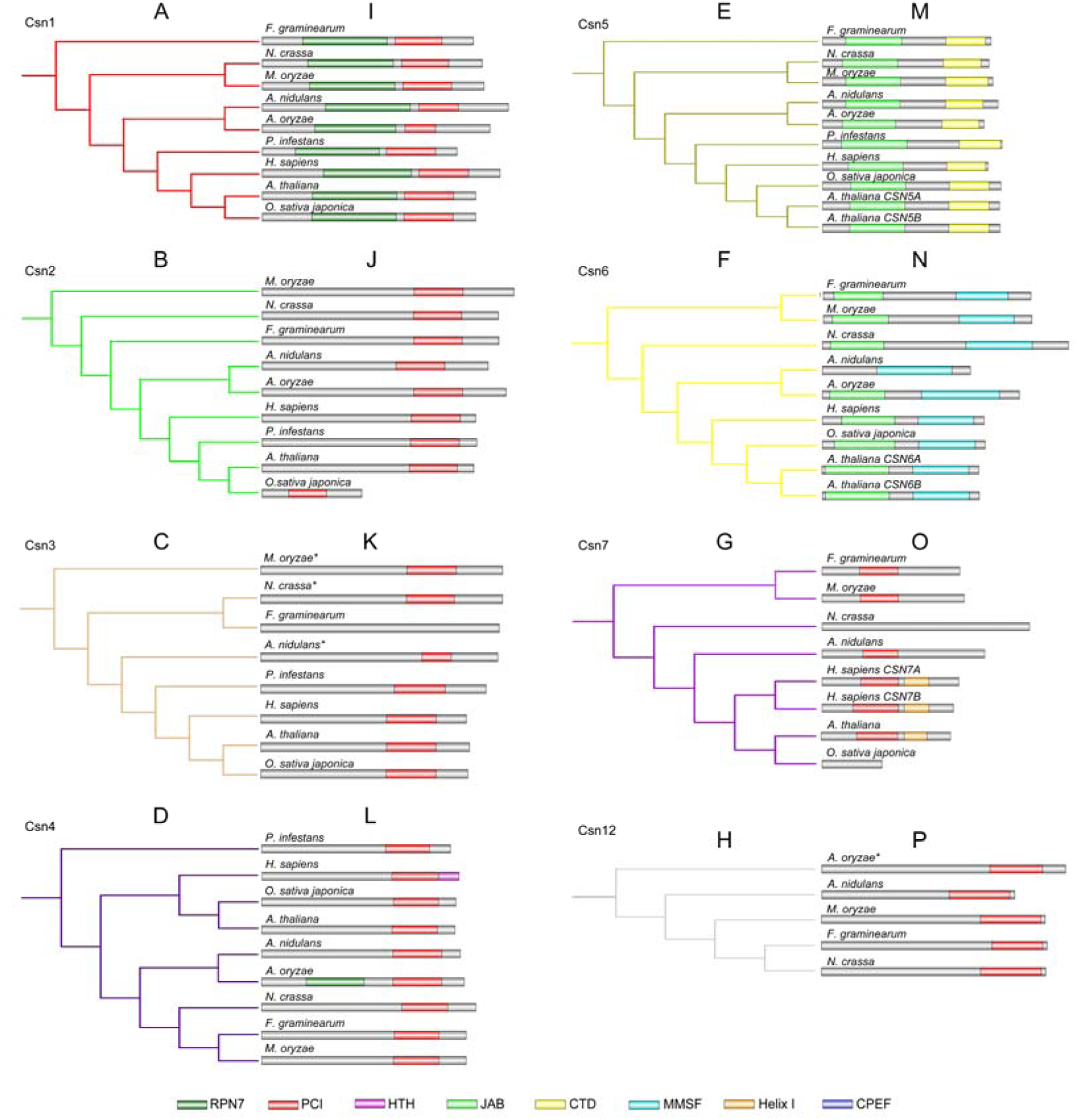
Phylogenetic and domain composition of putative orthologs of CSN subunits retrieved from different organisms. A-H, The cladogram represents the phylogenetic relationship between CSN complex subunits identified in selected organisms. The likelihood neighbor-joining trees were constructed with the MEGA-X software. I-P, The distribution and functional domain architecture of CSN subunits identified in the selected organisms. Domain prediction was performed using the Pfam server, and sequences with significant similarity to functional domains were used to construct domain models for the CSN subunits using GPS IBS 1.0.2 software.

Comparative analyses showed that the composition and domain architecture of Csn1, Csn2, Csn4, Csn5, and Csn6 are relatively conserved across species. However, compared to plants (*A. thaliana* and *O. sativa japonica)*, animals (*H. sapiens)*, and oomycetes (*P. infestans)*, the Csn3 sequence obtained from *M. oryzae* and other filamentous fungi, including *A. nidulans, A. oryzae*, *F. graminearum, M. oryzae*, and *N. crassa*, lacks the conserved Proteasome, COP9, and Initiation factor 3 domains. In addition, while Csn7 sequences retrieved from *A. nidulans*, *F. graminearum*, and *M. oryzae* contained a single PCI, Csn7 orthologs in *H. sapiens* and *A. thaliana* possessed a helix (CTD) in addition to the PCI domain (Figure 1I-P). From these observations, we inferred that the intra- and inter-species structural divergence between the corresponding orthologs of CSN subunits might reflect functional diversity between individual CSN subunit orthologs in different organisms.

### Nuclear export and import signals likely regulate the dynamic nucleocytoplasmic localization of the CSN complex

Results obtained from the screening of NLS/NES in CSN subunits identified in *M. oryzae*, identified a strong NES consensus motif signature in Csn3, Csn5, Csn7, and Csn12 (NetNES/NN-Score >0.7) and a relatively weak NES consensus motif signature in Csn6 (NetNES/NN-Score =0.6), while Csn7 was the only subunit with a consensus NLS motif signature in *M. oryzae* (Figure 2A-B).

**Figure 2.**
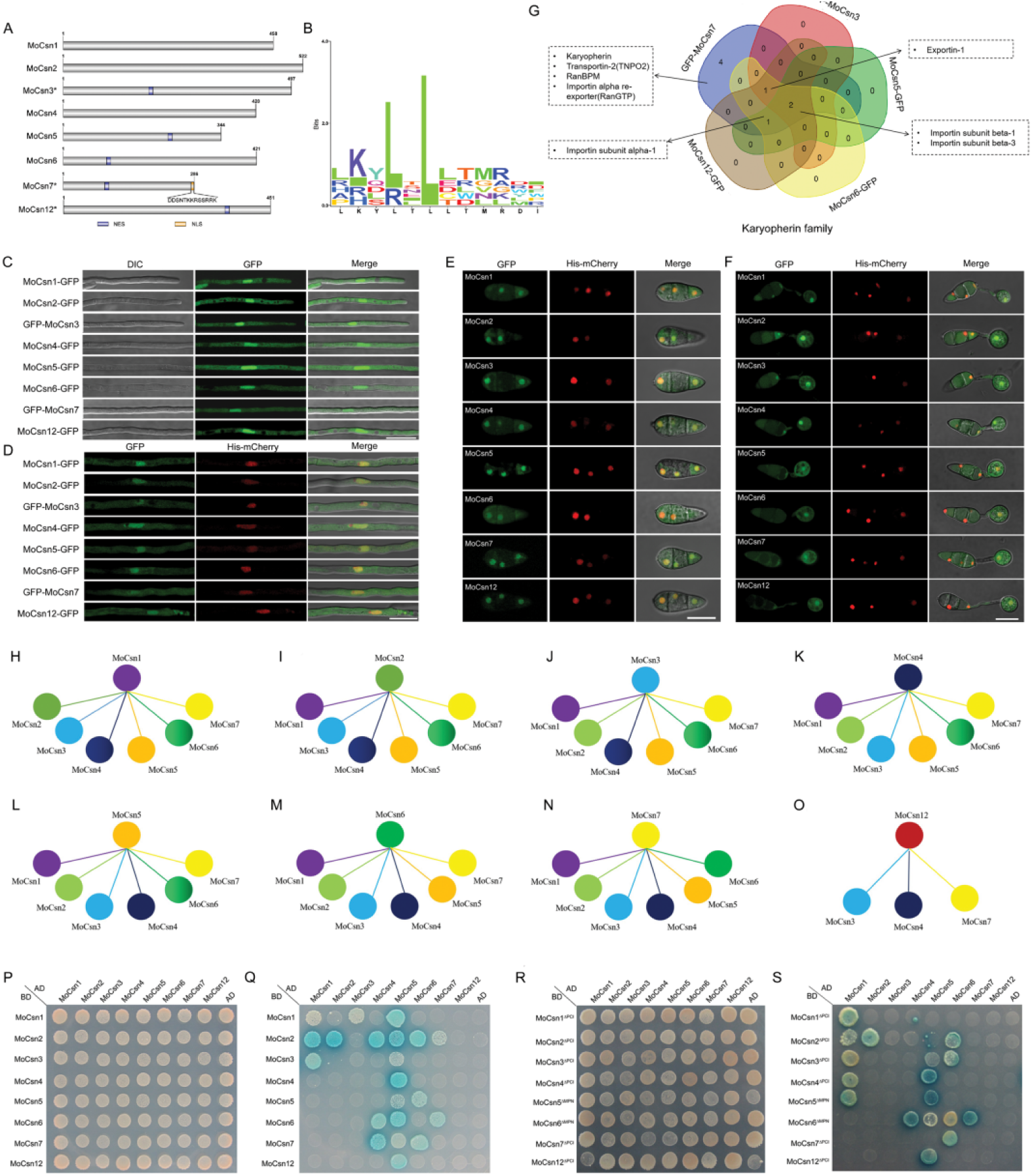
Detailed sequence feature analysis of individual CSN subunits and dynamic interaction network between CSN subunits and karyopherin protein family in *M. oryzae*. A, Schematic plot of NES and NLS signal peptide in the individual subunit of the *M. oryzae* CSN complex. B, Aligned sequence logo of NES signatures identified in MoCsn3, MoCsn5, MoCsn6, MoCsn7, and MoCsn12. C, Localization pattern of *M. oryzae* CSN subunits with GFP fused to their C-terminus (MoCsn-GFP) in vegetative hyphae of rice blast fungus. D-F, Co-localization of CSN subunits with His-mCherry in aberrant conidia and pathogenic development of the rice blast pathogen. G, Venn chart of proteins from the karyopherin family that interacts with the individual subunits of the Csn complex in *M. oryzae*. H-O, Interaction architecture prevailing between individual subunits of the MoCsn holocomplex based on mass spectrometry analysis. The interaction outlooks were constructed with co-immunoprecipitation data obtained from three technical replicates. Only subunits recovered from the immuno-complexes of the three replicates of the bait subunit are considered putative interactors. P, Yeast-two-hybrid assay showing the growth paired subunits of the CSN complex on synthetic defined minimal yeast media plates supplemented with Trp-Leu as a measure of interaction between the paired subunits MoCsn-AD and MoCsn-BD *in vitro*. Q, Growth of yeast transformants expressing the MoCsn-AD and MoCsn-BD on SD supplemented with Trp-Leu-His-Ade+α-gal. R, Yeast-two-hybrid assay showing the growth paired subunits of the CSN complex on SD minimal yeast media plates supplemented with Trp-Leu as a measure of interaction between the paired subunits MoCsn-AD and the domain deletion BD vector of MoCsn *in vitro*. S, Growth of yeast transformants expressing the MoCsn-AD and the domain deletion BD vector of MoCsn on SD supplemented with Trp-Leu-His-Ade+X-α-gal. Asterisks (*) represent *M. oryzae* CSN subunits with significantly high NES scores (≥0.7). Bar = 10 μm.

While most CSN subunits are exclusively localized in the nucleus in some cell types or organisms, recent work has shown that some subunits localize to both the nucleus and (Chamovitz and DENG, 1997; Wang et al., 2009). Therefore, to ascertain the localization and gain insights into the possible biological functions of CSN subunits in rice blast fungus, we monitored the subcellular localization of MoCsn subunits at different developmental stages in *M. oryzae* (hyphae, conidia, and appressorium) by fusing GFP fluorescent protein to the C-terminus of the eight CSN subunits consisting of the CSN complex in *M. oryzae*. Microscopy revealed that, except for MoCsn3-GFP and MoCsn7-GFP, the other subunits, i.e., MoCsn1-GFP, MoCsn2-GFP, MoCsn4-GFP, MoCsn5-GFP, MoCsn6-GFP, and MoCsn12-GFP, accumulated strongly in the nucleus and were weakly localized to the cytoplasm (Figure 2C and Figure S1A-B). Contrary to our expectations, MoCsn3-GFP and MoCsn7-GFP displayed persistent cytoplasmic localization patterns (Figure S1C-D).

Since the NLS signal peptide is located at the C-terminal of MoCsn7, we reasoned that fusing GFP to the C-terminal might result in the mislocalization of MoCsn7. To validate this possibility, we generated N-terminal GFP-fusion constructs for MoCsn3 and MoCsn7 (GFP-MoCsn3 and GFP-MoCsn7, respectively) and transformed them independently into wild-type protoplasts. Microscopy of individual positive transformants harboring GFP-MoCsn3 and GFP-MoCsn7 revealed strong GFP-MoCsn3 and GFP-MoCsn7 fluorescence signals in the nucleocytoplasmic regions.

However, GFP-MoCsn3 and GFP-MoCsn7 fluorescence signals in the cytoplasm were weaker than those recorded in the nucleus (Figure 2C and Figure S1A-B). From these observations, we inferred that the NLS region (C-terminal) is essential for the proper localization of MoCsn7. In addition, the C-terminus of MoCsn3 appears to be essential for its nuclear localization.

We also demonstrated that the individual CSN subunits identified in *M. oryzae* partially colocalized with H1-mCherry fluorescence (a nuclear marker) at all developmental stages, including mycelia, conidia, and appressorium formation (Figure 2D-F). The nucleocytoplasmic localization pattern observed for the eight CSN subunits identified in rice blast fungus suggests that the COP9-complex likely contributes to the development of *M. oryzae* via direct or indirect modulation of cellular and developmental processes in both the nucleoplasm and cytoplasm.

Furthermore, we screened for protein interactions using co-immunoprecipitation of GFP-MoCsn3, MoCsn5-GFP, MoCsn6-GFP, GFP-MoCsn7, and MoCsn12-GFP. Our data revealed that karyopherin (MGG_01449), transportin-2 (TNPO2/MGG_09208), RanBPM (MGG_00753), and importin alpha re-exporter (RanGTP/MGG_03994) were exclusively found in the GFP-MoCsn7 immuno-complex. GFP-MoCsn7, GFP-MoCsn3, and MoCsn12-GFP immunoprecipitated complexes contained exportin-1 (MGG_03994). We also found that importin subunit alpha (MGG_15072) interacted with MoCsn5-GFP, MoCsn6-GFP, MoCsn7, and MoCsn12-GFP. Meanwhile, importin subunit beta-1 (MGG_03668) and importin subunit beta-3 (MGG_03537) were present in the interaction complexes of all the CSN subunits tested (Figure 2G). Therefore, five subunits (MoCsn3, MoCsn5, MoCsn6, MoCsn7, and MoCsn12) may interact with exportins and play individual or overlapping roles in exporting other subunits and associated cargo from the nucleus to cytoplasm in *M. oryzae*.

Additional co-immunoprecipitation and yeast two-hybrid analysis results revealed dynamic physical interactions between MoCSN subunits in vivo (Figure 2H-O and Table S1) and *in vitro* (Figure 2P-Q). We also demonstrated that truncation of the PCI motif in MoCsn1 (MoCsn1^ΔPCI^), MoCsn2 (MoCsn2^ΔPCI^), and MoCsn7 (MoCsn7^ΔPCI^) abolished the physical interaction between MoCsn1^ΔPCI^-BD and MoCsn1-AD and MoCsn5-AD; MoCsn2^ΔPCI^ and MoCsn4-AD, MoCsn5-AD, and MoCsn7-AD; and MoCsn7^ΔPCI^-BD and MoCsn4-AD and MoCsn5-AD. Interestingly, truncation of the PCI motif in MoCsn4 triggered a physical interaction between MoCsn4^ΔPCI^-BD and MoCsn1-AD. Deletion of the MPN motif led to a physical interaction between MoCsn6^ΔMPN^-BD and MoCsn1-AD (Figure 2R-S). Based on these observations, we propose that the PCI and MPN domains, particularly the PCI domains in MoCsn1 and MoCsn2, play a crucial role in regulating the structural conformations required to facilitate the interaction network between CSN subunits in rice blast fungus.

### Targeted gene deletion of NES and NLS motif-containing CSN complex subunits suppressed morphological development and compromised cell wall integrity and stress tolerance in *M. oryzae*

Using homologous recombination, we successfully generated targeted gene deletion strains of *MoCSN7* and other subunits containing an NES signal motif, including *MoCSN3, MoCSN5, MoCSN6*, and *MoCSN12*. The targeted gene deletion strains generated for the five subunits selected for functional characterization analyses in this study, along with their respective complementation strains, were confirmed by Southern blotting assays (Figure S2A-E).

Comparative colony diameter analyses showed that while targeted gene replacement of *MoCSN3* and *MoCSN12* caused a substantial but non-significant reduction in the colony diameter of the Δ*Mocsn3* and Δ*Mocsn12* strains, targeted gene replacement of *MoCSN5*, *MoCSN6*, and *MoCSN7* caused a significant reduction in the vegetative development of the individual deletion strains, particularly in Δ*Mocsn7* (Figure 3A-B). We also observed hyper-melanization in the Δ*Mocsn3*, Δ*Mocsn5*, Δ*Mocsn6*, and Δ*Mocsn7* strains cultivated with solid and liquid CM (Figure 3C).

**Figure 3.**
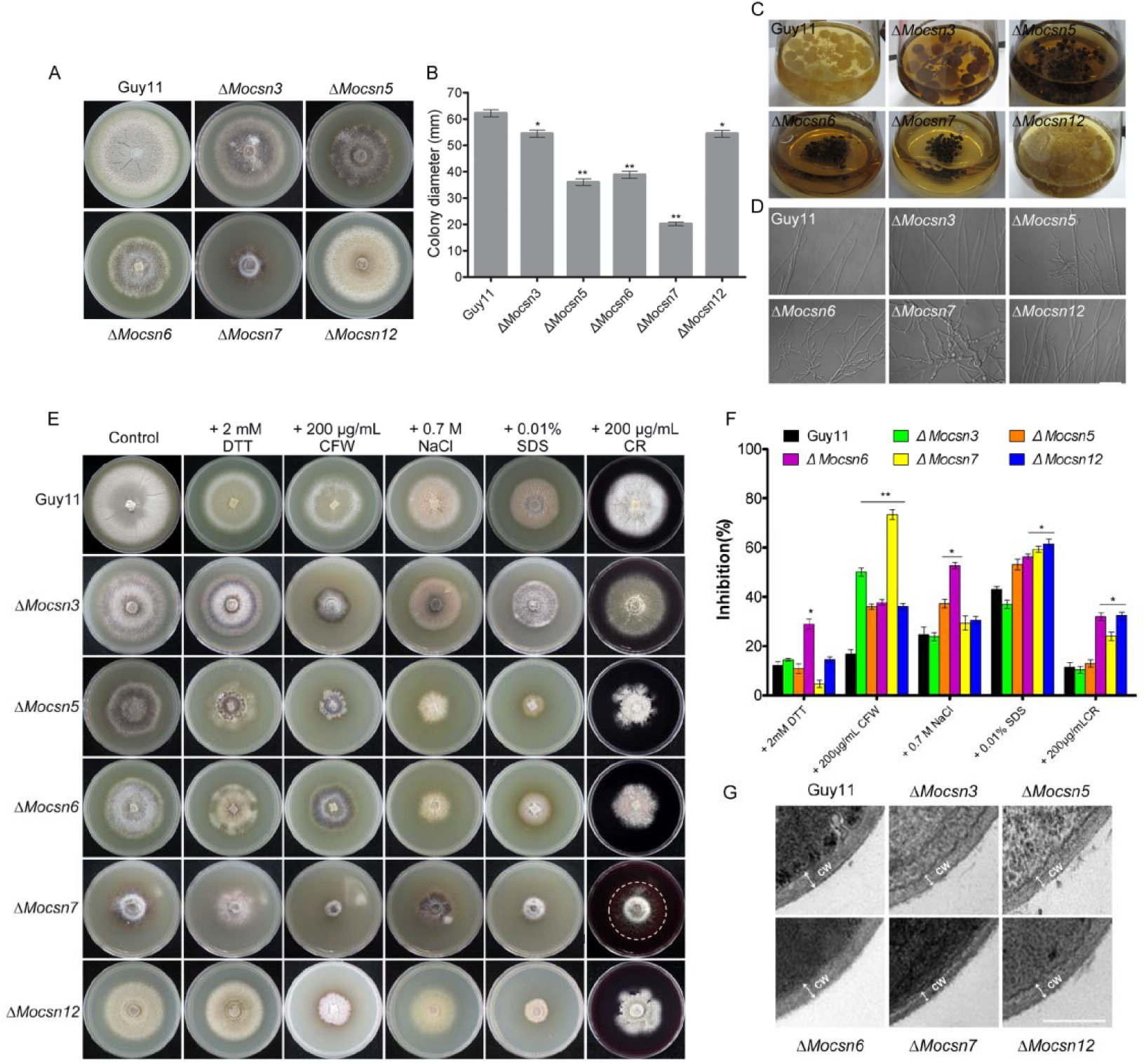
The impact of individual CSN subunit deletion on vegetative growth, hyphal branching, stress tolerance, cell membrane, and cell wall integrity in *M. oryzae*. A, Comparative vegetative growth and colony morphology of Δ*Mocsn3*, Δ*Mocsn5*, Δ*Mocsn6*, Δ*Mocsn7*, *ΔMocsn12*, and the strain grown on CM for 10 days. B, A statistical representation of average colony diameters for the individual strains grown on CM for 10 days. C, Hyper-melanization of the Δ*Mocsn3*, Δ*Mocsn5*, Δ*Mocsn6*, and Δ*Mocsn7* strains compared to that of the *ΔMocsn12* and the wild-type strains. Individual strains were cultured in liquid CM and incubated under optimum growth conditions for 3 days. D, Micrograph showing the impact of targeted gene deletion of *MoCSN3, MoCSN5, MoCSN6, MoCSN7*, and *MoCSN12* on hyphal branching in *M. oryzae.* Bar = 10 μm. E, The colony morphology of Δ*Mocsn3*, Δ*Mocsn5*, Δ*Mocsn6*, Δ*Mocsn7*, Δ*Mocsn12*, and the wild-type strains cultured on CM medium supplemented independently with 2 mM DTT, 200 μg/mL CFW, 0.7 M NaCl, 0.01% SDS and 200 μg/mL CR. F, A statistical representation of the inhibitory effects of individual stress-inducing osmolytes on the vegetative development of Δ*Mocsn3*, Δ*Mocsn5*, Δ*Mocsn6*, Δ*Mocsn7*, Δ*and Mocsn12* strains compared to the wild-type strains. G, Transmission electron microscopy images showing the morphological architecture of hyphae cells in wild-type strain compared to that of the individual gene deletion strains of the *MoCSN* complex. Bar =0.5 μm. Statistical analyses in “B” were performed with consistent results obtained from three biological replicates, each consisting of three technical replicates. Error bars represent the standard deviation between replicates, while single (“*”) and double (“**”) asterisks represent a statistically significant difference of (P≤ 0.05) and (P≤ 0.01), respectively.

Further microscopy-based morphological assessment of vegetative hyphae produced by individual gene deletion strains revealed that MoCsn5, MoCsn6, and MoCsn7 positively regulate hyphal morphogenesis in *M. oryzae.* Hence, targeted gene disruption of *MoCSN5*, *MoCSN6*, and particularly *MoCSN7* triggered hyper-hyphal branching and severely compromised the morphology of vegetative hyphae produced by the *ΔMocsn5*, *ΔMocsn6*, and *ΔMocsn7* strains (Figure 3D). Furthermore, reintroducing full-length coding sequences for individual *MoCSN* genes into their corresponding mutant strains fully rescued the vegetative growth defects observed in the individual *ΔMocsn* strains (Figure S3A-B). These results show that although the NLS and NES signal peptide motif-containing subunits of the CSN complex are not essential for the direct survival of rice blast fungus, MoCsn5, MoCsn6, and MoCsn7 play essential roles in the normal morphological differentiation and vegetative development of *M. oryzae*.

Sensitivity assessment of the individual strains to selected osmolytes showed that targeted deletion of *MoCSN3, MoCSN5, MoCSN6, MoCSN7*, and *MoCSN12* generally compromised the stress tolerance of *M. oryzae* and rendered defective strains particularly sensitive to CFW and SDS. In addition, we observed that only targeted disruption of the *MoCSN7* gene compromised cell wall integrity and resulted in leakage of cell wall-associated enzymes during the growth of the *ΔMocsn7* strains on CM supplemented with CR. We also observed that, except for *ΔMocsn6*, the other defective strains, particularly *ΔMocsn7*, exhibited profound immunity against ER-associated reductive stress-inducing agents (Figure 3E-F and Figure S4). These results indicate that individual subunits of the CSN complex exert different influences on reductive and oxidative stress tolerance in *M. oryzae*.

Based on these observations, we hypothesized that MoCsn7 plays a core role in promoting oxidative stress tolerance in rice blast fungi by regulating the activities of cell wall biogenesis enzymes. To validate this position, we observed the cell wall texture of individual strains under a transmission electron microscope. Transverse cross-sections of vegetative hyphae cells from the *MoCSN3, MoCSN5, MoCSN6, MoCSN7*, and *MoCSN12* defective strains and wild-type strains revealed the formation of a thinner cell wall in the Δ*Mocsn3*, Δ*Mocsn5*, and Δ*Mocsn7* strains compared to the texture of the cell wall observed in Δ*Mocsn6*, Δ*Mocsn12*, and the wild-type strains (Figure 3G). The attenuation in cell wall texture and the leakage of hydrolytic enzymes, as manifested in the circular watermark preceding the hyphae growth front of the Δ*Mocsn7* strains grown on CR, partly confirms that the MoCsn7 subunit plays a direct or indirect role in promoting cell wall integrity and oxidative or cell wall responsive stress tolerance in *M. oryzae*.

### Targeted disruption of genes coding for NLS/NES-containing subunits of the MoCSN complex differentially compromise sexual/sexual reproduction and pathogenicity/virulence in *M. oryzae*

Although previous studies have shown that CSN-dependent photomorphogenesis crucially regulates asexual sporulation in Neurospora(Zhou et al., 2012), the contribution of the CSN complex to reproductive development in plant pathogenic fungi, including globally destructive rice blast pathogens, has not been extensively examined. Conidiation assays revealed that while targeted gene replacement of *MoCSN3* and *MoCSN12* triggered a significant reduction in asexual reproduction, targeted disruption of *MoCSN5, MoCSN6*, and *MoCSN7* completely abolished asexual sporulation (Figure 4A-B). However, we observed that the individual subunits of the MoCSN complex investigated in this study were indispensable for conidiophore genesis in rice blast fungus (Figure 4C). We conclude that NLS/NES motif-containing subunits (and, by extension, the CSN complex) likely promote asexual sporulation in *M. oryzae* by modulating pathways or processes independent of conidiophore formation.

**Figure 4.**
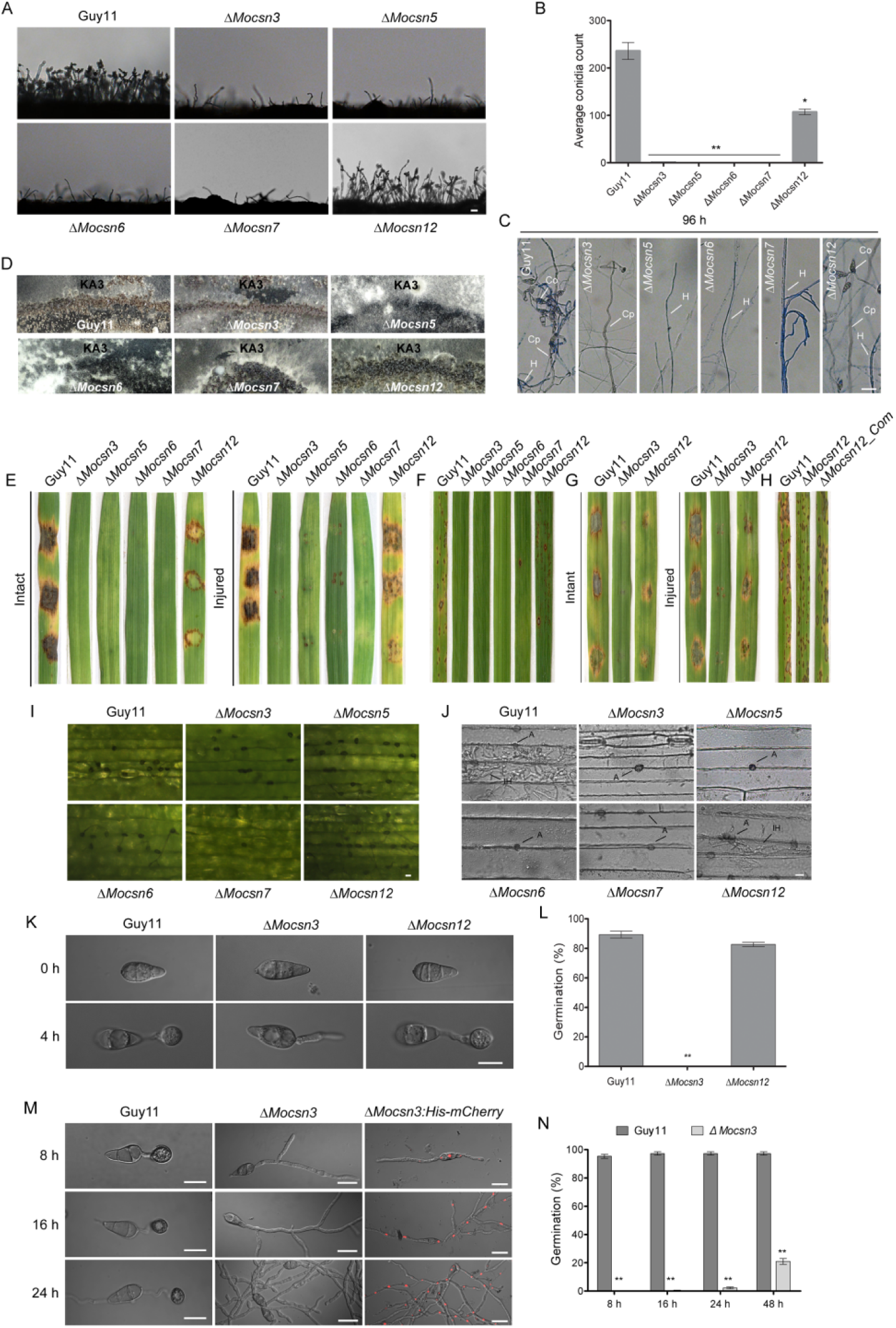
Targeted gene disruption of *MoCSN* subunits attenuated sporulation, pathogenicity, virulence, and pathogenic differentiation in rice blast fungus. A, The micrograph represents a comparative microscopic assessment of conidiophore formation upon deleting individual *MoCSN* subunits and the wild-type strain exposed to light 24 h after the vegetative growth phase. Bar =20 μm. B, Statistical analysis of conidiation in individual CSN subunit complementation strains. C, Examination of conidiophore formation in the Δ*Mocsn3*, Δ*Mocsn5*, Δ*Mocsn6*, Δ*Mocsn7, and* Δ*Mocsn12* strains compared to wild-type strains at 96 h after induction of conidiogenesis. The strains were treated with lactophenol cotton blue solution to differentiate conidiophores from vegetative hyphae and conidia. Bar =20 μm. D, Sexual sporulation in strains defective for CSN genes crossed with compatible mating-type strain KA3 mating-inducing culture media for 3 weeks. E, Hyphae-mediated infection characteristics of individual defective strains on intact and injured leaves excised from barley seedlings 7 days post-inoculation (dpi). F, Infection results were obtained from the inoculation of 3-week-old CO39 rice cultivars with mycelia suspensions prepared with mycelium from the individual defective strains 7 dpi. (G-H) Conidia-mediated infection characteristics of Δ*Mocsn3, ΔMocsn12*, and the wild-type strains against barley and CO39 rice cultivars. Excised barley leaves were drop-inoculated with 20 μL spore suspension, while the CO39 rice seedlings were spray-inoculated with the spore suspensions. I, Comparative formation of appressorium-like structures in the *MoCSN* defective strains and the wild-type on inoculated barely leaves after 3 dpi. J, The histopathogenicity assay shows penetration characteristics of the targeted gene deletion strains generated for individual subunits of the *MoCSN* complex inoculated on barley leaves for 48 h compared to the wild-type. “A and IH” represent appressorium, and invasive hyphae, respectively. K, The micrograph portrays conidia germination and appressorium formation in the Δ*Mocsn3* and *ΔMocsn12* strains inoculated on hydrophobic slides 4 hpi compared to the wild-type strains. White Bar = 10 μm magnification. L, Statistical presentation of results obtained from conidia germination and appressorium formation in the Δ*Mocsn3, ΔMocsn12*, and the wild-type strains observed and counted under a light microscope. M, Delayed formation and hyper-germ tube branching in Δ*Mocsn3* strains inoculated on hydrophobic slides after extended periods of 8, 16, and 24 hpi. Bar = 10 μm. N, Statistical presentation of results obtained from appressorium formation in the Δ*Mocsn3 strain* after extended periods of 8, 16, and 24 hpi on hydrophobic slides compared to the wild-type strains. Statistical analyses were performed using results from three independent biological experiments, with each consisting of three technical replicates. The error represents standard deviation, while single “*” and double “**” asterisks represent a significant difference of (P≤ 0.05) and (P≤ 0.01), respectively.

Additionally, we evaluated sexual sporulation in the *ΔMocsn3*, Δ*Mocsn5*, Δ*Mocsn6*, Δ*Mocsn7*, and Δ*Mocsn12* strains compared to the wild-type strain using pairwise inoculation of the individual strains with the mating-type KA3 strain on oat agar (OA) culture media. Somatotype microscopy examination of the mating compatibility of the individual strains with KA3 strains revealed the formation of perithecia at the intersection zones in *ΔMocsn3* and KA3, *ΔMocsn5* and KA3, *ΔMocsn7* and KA3, and *ΔMocsn12* and KA3, indicating mating compatibility between the respective defective strains and the KA3 strain. However, the Δ*Mocsn6* and KA3 strains failed to form perithecia at convergence (Figure 4D). These results indicate that MoCsn5, MoCsn6, and MoCsn7 likely play overlapping roles in promoting the asexual development of rice blast fungus. We also speculate that MoCsn6 likely forms a complex with other CSN subunits to modulate the progression of sexual development in *M. oryzae*.

We showed that, except for *MoCSN12*, targeted gene disruption of *MoCSN3* caused a 98% reduction in asexual sporulation in Δ*Mocsn3* strains, whereas Δ*Mocsn5*, Δ*Mocsn6*, and Δ*Mocsn7* strains completely lost their asexual sporogenesis abilities. Therefore, to assess the significance of the individual *MoCSN* genes in the pathogenicity and virulence of *M. oryzae*, mycelia obtained from the wild-type, *ΔMocsn3, ΔMocsn5, ΔMocsn6, ΔMocsn7, ΔMocsn12*, and complementation strains cultured in CM for 3 days were used to inoculate the leaves of 7-day old blast-susceptible barley (Golden promise cultivar) seedlings. At the same time, 14-day-old seedlings of the homogeneous blast-susceptible CO39 rice cultivar were spray-inoculated with suspensions of mycelia obtained from individual strains. We observed that while *ΔMocsn3, ΔMocsn5, ΔMocsn6*, and *ΔMocsn7* failed to initiate hyphae-mediated blast infection on “intact” and injured leaves of barley and rice seedlings, the *ΔMocsn12* strain caused blast lesions in both barley and rice seedlings. However, the blast symptoms caused by the Δ*Mocsn12* strain were less severe than those of the wild-type and complementation strains (Figure 4E-F). Conidia harvested from the Δ*Mocsn3*, *ΔMocsn12*, and wild-type strains were used for spore suspension drop-inoculation (*ΔMocsn3, ΔMocsn12*, and the wild-type) and spray (Δ*Mocsn12* and Guy11) inoculation of susceptible CO39 rice seedlings. Records from conidia-mediated infection assays confirmed that targeted deletion of *MoCSN3* rendered the Δ*Mocsn3* strains non-pathogenic, while *MoCSN12* deletion suppressed the virulence of rice blast fungus (Figure 4G-H).

To unravel the possible link between subunits of the CSN complex and the pathogenesis of *M. oryzae*, we monitored pathogenic differentiation in the individual defective strains by comparatively examining the formation of hyphal-tip appressorium-like structures in the *ΔMocsn3, ΔMocsn5, ΔMocsn6, and ΔMocsn7* strains inoculated on barley leaves. We also assayed conidia-associated appressorium formation in conidia obtained from *ΔMocsn3, ΔMocsn12*, and wild-type strains on appressorium-inducing hydrophobic coverslips. The results of these investigations revealed that targeted gene replacement of *MoCSN3*, *MoCSN5*, *MoCSN6*, *MoCSN7*, and *MoCSN12* had no visible adverse effects on the formation and maturation of hyphae-tip appressorium-like structures in *M. oryzae* (Figure 4I). Although histopathological bioassays showed that the Δ*Mocsn3*, Δ*Mocsn5*, Δ*Mocsn6*, and Δ*Mocsn7* strains could form fully melanized hyphae-tip appressorium-like structures during pathogen-host interaction (PHI), these appressorium-like structures could not differentiate into penetration pegs and were unable to invade the epidermal tissues of the host plant. The Δ*Mocsn12* strain, on the other hand, produced functional penetration pegs and actively invaded the host cells. However, the tissue colonization efficiency (virulence) of Δ*Mocsn12* was drastically reduced compared to that of the wild type (Figure 4J).

Furthermore, we demonstrated that the *MoCSN12* deletion had no adverse effects on conidial germination and appressoria morphogenesis in the Δ*Mocsn12* strain (Figure 4K-L). However, targeted disruption of *MoCSN3* delayed the onset of conidia germination, compromised the polarized development of germ tubes, triggered hyper-branching of germ tubes, and severely suppressed appressorium formation in the Δ*Mocsn3* strain (Figure 4M-N). From these results, we concluded that the NLS/NES domain-containing subunits of the CSN complex promote the pathogenicity and virulence of rice blast fungus, possibly by regulating multiple pathways with the pathogenic development of filamentous fungal pathogenes.

### MoCsn6 and MoCsn7 are essential for photo-dependent ubiquitin homeostasis in ***M. oryzae***

Given the profound upregulation in the transcription levels of genes linked to the SCF pathway, we determined whether NLS/NES motif-containing subunits of the CSN complex physically interact with components of the SCF complex. First, we searched for SCF complex-associated proteins in the interactome library generated. Of the 23 SCF complex-associated proteins recovered from the co-immunoprecipitation complexes of the individual subunits, 2/26, 5/26, and 1/26 co-immunoprecipitated exclusively with MoCsn6-GFP, GFP-MoCsn7, and MoCsn12-GFP, respectively. Furthermore, we observed that 5/26, 7/26, 1/26, 1/26, and 1/26 ubiquitin pathway-associated proteins were exclusively present in the immuno-complexes of GFP-MoCsn3 and MoCsn5-GFP; MoCsn6-GFP and MoCsn12-GFP; GFP-MoCsn3, MoCsn5-GFP, and MoCsn6-GFP; GFP-MoCsn3, MoCsn5-GFP, MoCsn6-GFP, and GFP-MoCsn7; and GFP-MoCsn3, MoCsn5-GFP, MoCsn6-GFP, and GFP-MoCsn3, respectively. In addition, 3/26 SCF-associated proteins were recovered as common interaction proteins of GFP-MoCsn3, MoCsn5-GFP, MoCsn6-GFP, GFP-MoCsn7, and MoCsn12-GFP (Figure 5A). These results suggest that, among the five CSN subunits investigated in this study, MoCsn7 potentially assumes a more extensive regulatory role in the progression of protein ubiquitination and degradation events in *M. oryzae*.

**Figure 5.**
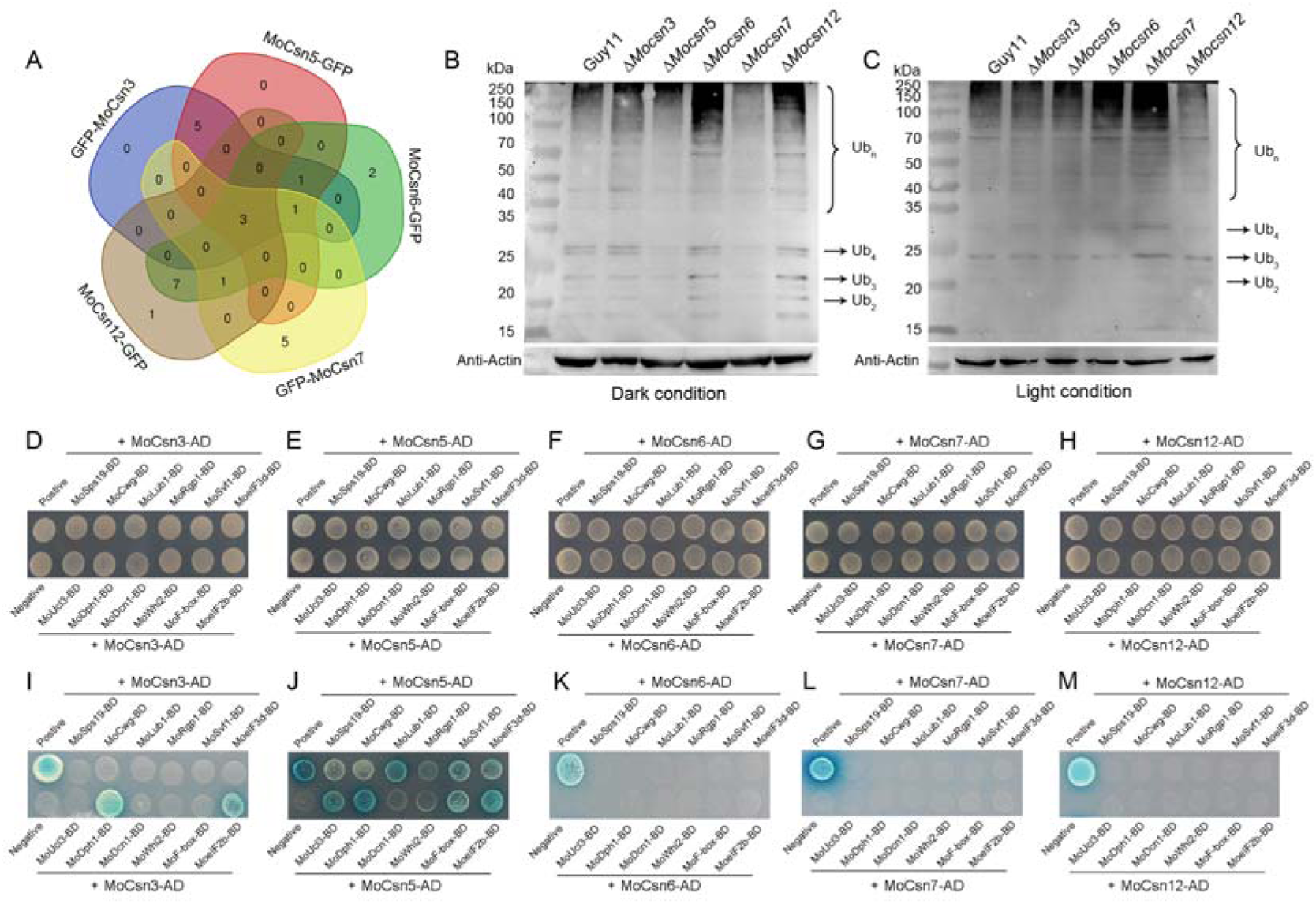
The interaction network between NLS/NES motif containing CSN subunits and SCF pathway-associated proteins and the impact of NLS/NES motif containing dysfunctioning CSN subunits on ubiquitination processes in *M. oryzae.* A, The Venn chart showing the number of SCF-associated proteins recovered from immuno-complexes of the individual NLS/NES motif-containing subunits of the MoCSN complex. B-C, Western-blot image showing ubiquitination levels in the Δ*Mocsn3*, Δ*Mocsn5*, Δ*Mocsn6*, Δ*Mocsn7*, Δ*and Mocsn12* strains compared to the wild-type strains in response to dark and light treatment. D-H, Yeast-two-hybrid assay showing the growth of NLS/NES motif containing CSN subunits paired with SCF pathway-associated proteins on synthetic defined minimal yeast media plates supplemented with Trp-Leu to monitor the likely existence of physical interaction between the paired subunits MoCsn-AD and MoSCF-BD *in vitro.* I-M, Yeast-two-hybrid assay showing the growth of NLS/NES motif containing CSN subunits paired with SCF pathway-associated proteins on synthetic defined minimal yeast media plates supplemented with Trp-Leu-His-Ade+X-α-gal to monitor the likely existence of physical interaction between the paired subunits MoCsn-AD and MoSCF-BD *in vitro*.

Generally, the COP9-signalosome complex facilitates the termination of ubiquitination by decoupling neddylases from the Cullen protein(Suisse *et al*., 2018). Therefore, total proteins were extracted from the Δ*Mocsn3*, Δ*Mocsn5*, Δ*Mocsn6*, Δ*Mocsn7*, Δ*Mocsn12*, and the wild-type strains cultured independently under continuous light and dark conditions for 3 days were used for a comparative examination of poly-ubiquitination levels in the individual strains by performing anti-ubiquitin antibody blotting assays. This provided further insight into the direct impacts of *MoCSN3*, *MoCSN5*, *MoCSN6*, *MoCSN7*, and *MoCSN12* gene deletions on the progression of the ubiquitination process in rice blast fungus, the possible link between CSN dysfunction, and the observed upregulation in the expression pattern of genes coding for SCF pathway-associated proteins, especially in the Δ*Mocsn6* and Δ*Mocsn7* strains. The results obtained from the ubiquitination assessment assays revealed a substantial decrease in the accumulation of ubiquitin in the Δ*Mocsn7* and Δ*Mocsn5* strains under dark conditions compared to the remarkable accumulation of ubiquitin in the wild-type and the other defective strains, particularly the Δ*Mocsn6* strain under the same culture conditions (Figure 5B). Interestingly, the accumulation of ubiquitin in the Δ*Mocsn7* strain was substantially enhanced and higher than that observed in the wild-type, Δ*Mocsn3*, Δ*Mocsn5*, Δ*Mocsn6*, and Δ*Mocsn12* strains under continuous light conditions (Figure 5C). Based on these observations, we reasoned that MoCsn6 likely acts in tandem or combined with other subunits to facilitate ubiquitination under photo-deficient conditions. The MoCsn7 subunit promotes the progression of photogenic ubiquitination in *M. oryzae*.

Subsequently, we performed RT-qPCR assays to validate the expression of genes coding for ubiquitination pathway-associated proteins. This included deubiquitination-protection protein dph1(MoDPH1), ubiquitin homeostasis protein lub1(MoLUB1), defective in Cullin neddylation protein 1 (MoDCN1), ubiquitin C-terminal hydrolase L3 (MoUCL3), ubiquitin-conjugating enzyme (MoUCE), E3 ubiquitin-protein ligase ptr1 + RNA transporter 1 (MoPTR1), ubiquitin carboxyl-terminal hydrolase 6 (MoUCL6), SCF E3 ubiquitin ligase complex F-box protein grrA (MoGRRA), and ESCRT-I component (MoESCRT1); genes coding for putative interacting karyopherin proteins (RanBPM/MoBPM, transportin-2/MoTSP2, exportin-1/MoEXP1, and ranGTPase-activating protein 1/MoRGP1); and genes coding for putative interactors we perceived to directly, or indirectly associate with the physiological and pathological defects observed in the defective strains, including eukaryotic translation initiation factor3 subunit D (MoeIF3D), survival factor protein 1 (MoSVF1), translation initiation factor eIF-2B subunit alpha (MoeIF2B), UV excision repair protein Rad23 (MoRad23), stress-induced phosphoprotein 1 (MoSIP1), sporulation protein SPS19 (MoSPS19), general stress response protein Whi2 (MoWHI2), and cell wall glucanosyltransferase (MoCWG) recovered from the immuno-complex of the individual CSN subunits investigated in this study. Compared to *MoCSN3, MoCSN5, MoCSN6*, and *MoCSN12* we observed that targeted gene replacement of *MoCSN7* significantly inhibited the expression of genes coding for *MoDPH1, MoLUB1, MoDCN1, MoBPM, MoUCL3, MoPTR1, MoSIP1, MoSPS19*, and *MoCWG* (Figure S5A-B). In addition, yeast two-hybrid bioassays revealed direct interactions between MoCsn3-AD and MoDph1-BD, MoCsn3-AD and MoeIf2b-BD, MoCsn5-AD and MoUcl3-BD, MoCsn5-AD and MoDph1-BD, MoCsn5-AD and MoLub1-BD, and MoCsn5-AD and MoeIf2b-BD. Interestingly, none of the selected proteins directly interacted with MoCsn7 *in vitro* (Figure 5F-J). We hypothesized that the regulation of downstream substrates of the CSN complex, including MoDPH1, MoLub1, MoDcn1, MoBpm, MoUcl3, MoPtr1, Moip1, MoSps19, and MoCwg, by the MoCsn7 subunit is likely not mediated by physical interaction. Hence, targeted gene disruption of *MoCSN7* attenuates the transduction of upstream signals required for optimum transcription.

To validate this hypothesis, MoLub1, MoDcn1, and MoSps19 were independently expressed in the Δ*Mocsn3*, Δ*Mocsn*5, and Δ*Mocsn*7 strains under the constitutive ribosomal protein 27 (RP27) promoter(Dong et al., 2009). Results obtained from the phenotypic evaluation of individual strains harboring the MoDcn1, MoLub1, and MoSps19 overexpression constructs showed that constitutive expression of MoDcn1, MoLub1, and MoSps19 resulted in the substantial but differential restoration of growth and conidiation defects associated with MoCsn3, MoCsn5, MoCsn6, MoCsn7, and MoCsn12 defective strains. Overexpression of MoDcn1 and MoSps19 also abolished hyper-melanization exclusively in the Δ*Mocsn7* strains. Meanwhile, overexpression of MoDcn1 in the wild-type strain triggered a remarkable reduction in the conidiation characteristics of the Guy11-MoDcn1-OE strains (Figure S5C-E). Finally, the constitutive expression of MoDcn1 exclusively rescued the pathogenicity defects in the Δ*Mocsn3* and Δ*Mocsn5* strains (Figure S5F). Therefore, we conclude that the CSN complex subunits, particularly MoCsn7, positively promote morphogenesis and conidiogenesis in rice blast fungus, partly through transcriptional or post-translational regulation of downstream genes coding for substrate proteins associated with the SCF pathway, including *MoDCN1, MoLUB1*, and *MoSPS19*. We further suggest that MoDcn1 may be under internal feedback regulation in rice blast fungi for optimum conidiogenesis.

### Comparative non-targeted metabolomic assessment of CSN dysfunction on **metabolomic differentiation in *M. oryzae***

With the fundamental knowledge that light-responsive reactions have a far-reaching influence on the progression of metabolic and cellular processes, we evaluated the impact of targeted gene replacement of selected NLS/NES motif-containing subunits of the CSN signalosome complex on metabolomic differentiation in rice blast fungus. Total crude extracts were obtained from the mycelia of the Δ*Mocsn3* strains (representing a reduction in conidiation exclusively), Δ*Mocsn5* strains (representing multiple phenotypic defects, including loss of pathogenicity, sporulation, photoperiodism, and growth), and Δ*Mocsn7* strains (to exclusively represent strains with defects in stress tolerance). Extracts obtained from the individual strains were used for QTOF-UPHPLC-assisted non-targeted metabolomic profiling(Norvienyeku et al., 2021). In total, 139, 408, 204, and 199 metabolites were exclusively recorded in the Δ*Mocsn3*, Δ*Mocsn5*, Δ*Mocsn7*, and Guy11 strains, respectively. Δ*Mocsn3*, Δ*Mocsn5*, and Δ*Mocsn7* shared 48 metabolites, while Δ*Mocsn3*, Δ*Mocsn5*, Δ*Mocsn7*, and Guy11 strains had 1364 metabolites in common (Table S2). Further principal component analyses (PCA) showed that the recorded metabolites were consistently present in at least five of the six independent repeats (Figure S6A-B) and confirmed the reliability of the total mycelium metabolome data generated for the individual strains in this study.

### Targeted gene deletion of *MoCSN3*, *MoCSN5*, and *MoCSN7* suppressed the generation of metabolites associated with key catabolism and metabolism pathways in *M. oryzae*

Membrane-associated saturated fatty acids significantly contribute to enforcing membrane integrity and protecting cells against leakage(Oberhauser et al., 2020). In *Saccharomyces cerevisiae*, disruption of CSN5 caused a significant reduction in ergosterol levels and severely accelerated the membrane’s permeability to monovalent and divalent cations(Licursi et al., 2014).

Comparative non-targeted metabolomic analyses showed that targeted gene deletion of *MoCSN3*, *MoCSN5*, and *MoCSN7* triggered an imbalance in the generation and enrichment of metabolites associated with essential fatty acid metabolism pathways in rice blast fungus. We observed that targeted gene disruption of *MoCSN3*, *MoCSN5*, and *MoCSN7* triggered the upregulation of metabolites associated with linoleic acid and sphingolipids. However, metabolites associated with arachidonic acid, saturated fatty acid, purine, secondary metabolite biosynthesis, and amino acid metabolic pathways were downregulated in the metabolomes of the Δ*Mocsn3*, Δ*Mocsn5*, and Δ*Mocsn7* strains. In addition, the generation of metabolites associated with pyrimidine, oxidative phosphorylation, and the TCA-cycle pathways was suppressed exclusively in the Δ*Mocsn7* strains (Figure S7A-F). Furthermore, intra-mutant strain pathway enrichment analyses showed relatively higher enrichment of metabolites associated with amino acids (arginine, alanine, aspartate, glutamate, proline, lysine, threonine, serine, glycine, phenylalanine, and tyrosine) in the metabolome of the Δ*Mocsn7* strains compared to other CSN defective strains (Figure S7G-L). From these observations, we propose that the CSN signalosome complex positively regulates the cross-talk between diverse metabolic and catabolic pathways in *M. oryzae*.

### The addition of metabolites down-regulated in MoCSN defective strains can partially rescue conidiation defects associated with *ΔMocsn* strains

Differential metabolomic analyses revealed that targeted gene replacement of *MoCSN3*, *MoCSN5*, and *MoCSN7* triggered a significant reduction in the enrichment of fatty acids and fatty acid derivatives, including eicosapentaenoic acid, prostaglandin b1, tretinoin, choline, cAMP, and succinate. In addition, the arachidonic acid levels were significantly inhibited in the metabolomes of Δ*Mocsn3* and Δ*Mocsn5* (Figure S7A-L). Additionally, we observed that interference in the CSN signalosome complex significantly suppressed the abundance of cytidine, uridine, adenosine monophosphate (AMP), and xanthine/hypoxanthine (derivatives of nucleotide biosynthesis and pyrimidine degradation pathways) in rice blast fungus (Figure 6A-B). Further targeted metabolomic analyses confirmed a substantial decrease in intracellular levels of cAMP and pyridoxamine. However, the intracellular levels of arachidonic acid, xanthurenic acid, and azatadine were higher in the Δ*Mocsn7* strain than in the Δ*Mocsn3*, Δ*Mocsn5*, Δ*Mocsn6*, Δ*Mocsn12*, and wild-type strains (Figure 6C-N)

**Figure 6.**
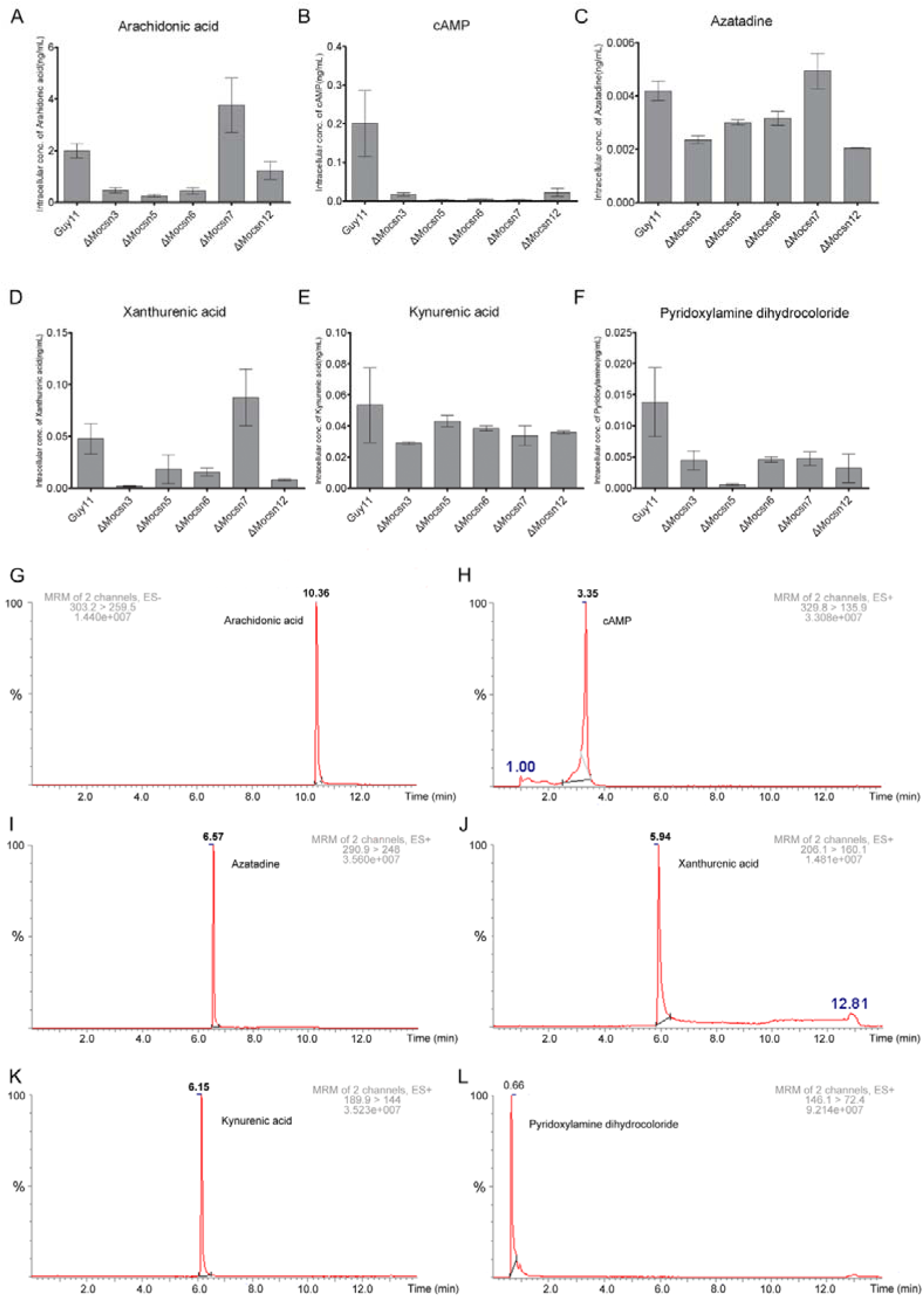
Targeted gene replacement of selected CSN subunits caused metabolome imbalance and severely suppressed the generation of cAMP in the Δ*Mocsn* strains. A, The heat-map cluster represents the abundance and intensity of metabolites in the Δ*Mocsn3*, Δ*Mocsn5*, and Δ*Mocsn7* strains compared to the wild-type strains collected in the positive (Pos.+) ionization mode. B, The heat-map clusters representing the abundance intensity of metabolites recorded in the Δ*Mocsn3*, Δ*Mocsn5*, and Δ*Mocsn7* strains compared to the wild-type strains collected in the negative (Neg.-) ionization mode. C-N, Results from chromatographic quantification of selected and deferentially expressed signaling molecules in the Δ*Mocsn3*, Δ*Mocsn5*, Δ*Mocsn6*, Δ*Mocsn7*, Δ*Mocsn12*, and wild-type strains. Comparative quantitative metabolomics were analyzed using ANOVA. The relative levels of the selected signaling molecules in the individual strains were measured using HPLC analytical standards as references. Differentially present metabolites with relative standard deviation (RSD) ˂30% t-test P-value (q-value) ˂0.05, mass error ≤± 3 with t-test P-value (q-value) ≤0.05, and computed relative standard deviation (RSD) ˂30%.

To ascertain whether the observed suppression in the generation of these metabolites is the limiting factor for growth, conidiation, stress tolerance, and pathogenic defects observed in the individual CSN subunit-defective strains, we assessed growth and conidiation in the Δ*Mocsn3*, Δ*Mocsn5*, Δ*Mocsn6*, Δ*Mocsn7*, and Δ*Mocsn12* strains cultured on CM supplemented independently with compounds that were differentially downregulated in the Δ*Mocsn3*, Δ*Mocsn5*, and Δ*Mocsn7* strains. The compounds used included arachidonic acid, cytidine, cAMP, xanthan, choline, uridine, kynurenine, and compounds that were exclusively present in the hyphae metabolome of the Δ*Mocsn7* strains, including quinoline, hydroxycinnamic acid, uric acid, 5-hydroxytryptamine, hydrochlorothiazide, glimepiride, milrinone, Rfaffinose, phosphoric acid, 4-hydroxy-3-methoxybenzyl alcohol, 5’-guanylic acid, equol, deoxycytidine, 5-hydroxytryptamine, nitrocatechol, mandelic acid, uridine monophosphate, quabbin, and phthalic acid. The results obtained from these bioassays showed that exogenous inclusion of 4-hydroxy-3-methoxybenzyl alcohol, 5’-guanylic acid, and equol caused a reduction in the vegetative growth of the wild-type strain and further aggravated the growth defects associated with the Δ*Mocsn6*, Δ*Mocsn7*, and Δ*Mocsn12* strains. Milrinone and quinoline exclusively restored conidiation defects associated with the Δ*Mocsn3*, Δ*Mocsn5*, and Δ*Mocsn6* strains. Deoxycytidine, uric acid, nitrocatechol, UMP, phthalic acid, and glimepiride selectively restored conidiation in the Δ*Mocsn3*, Δ*Mocsn5*, and Δ*Mocsn6* strains. Interestingly, these compounds failed to reverse the conidiation defects in the Δ*Mocsn7* strain (Table S3). In addition, we observed that saturated fatty acids and compounds upstream of metabolism and catabolism pathways, including arachidonic acid, linoleic acid, prostaglandin, oleic acid, pyrimidine, and choline, failed to rescue growth and conidiation defects associated with individual CSN gene deletion strains. Accordingly, we concluded that the CSN signalosome complex positively affects conidiogenesis in filamentous fungi through the synergistic regulation of diverse metabolic pathways. Furthermore, we reasoned that the physical presence of the *MoCSN7* subunit is essential for the biosynthesis of fatty acids and pyrimidine derivatives required to drive conidiation in *M. oryzae*.

### *MoCSN7* essentially promotes autophagic flux in *M. oryzae* by modulating the intracellular levels of cAMP under starvation

To examine the impact of targeted gene disruption of *MoCSN3, MoCSN5, MoCSN6, MoCSN7*, and *MoCSN12* on autophagosome formation, maturation, autophagolysosomal fusion, and degradation (autophagic influx), we performed a qPCR assay to monitor the expression pattern of genes coding for autophagy-related proteins (listed in Table S4) in the Δ*Mocsn3*, Δ*Mocsn5*, Δ*Mocsn6*, Δ*Mocsn7*, and Δ*Mocsn12* strains and compared it to that of the wild-type. Results obtained from transcriptomic analyses showed that targeted disruption of *MoCSN5, MoCSN6*, and *MoCSN7* significantly suppressed the expression of 24/24 autophagy-related genes identified in rice blast fungus. Deletion of *MoCSN3* and *MoCSN7* exclusively and significantly enhanced the expression of *MoATG4*, *MoATG3*, *MoATG6, MoATG7, MoATG10, MoATG15, MoATG16, MoATG17, MoATG26, MoATG27*, and *MoATG28*) respectively. Additionally, we observed significant upregulation in the expression of genes encoding *MoATG2*, *MoATG11*, *MoATG13*, and *MoATG4* in the Δ*Mocsn3* and Δ*Mocsn12* strains (Figure 7A). These results indicate that subunits of the CSN complex, particularly *MoCSN5*, *MoCSN6*, and *MoCSN7*, significantly influence the initiation and sequential progression of autophagic influx (autophagosome formation, maturation, autophagosome-vacuole/lysosome fusion, and likely cargo selection)(Klionsky and Eskelinen, 2014).

**Figure 7.**
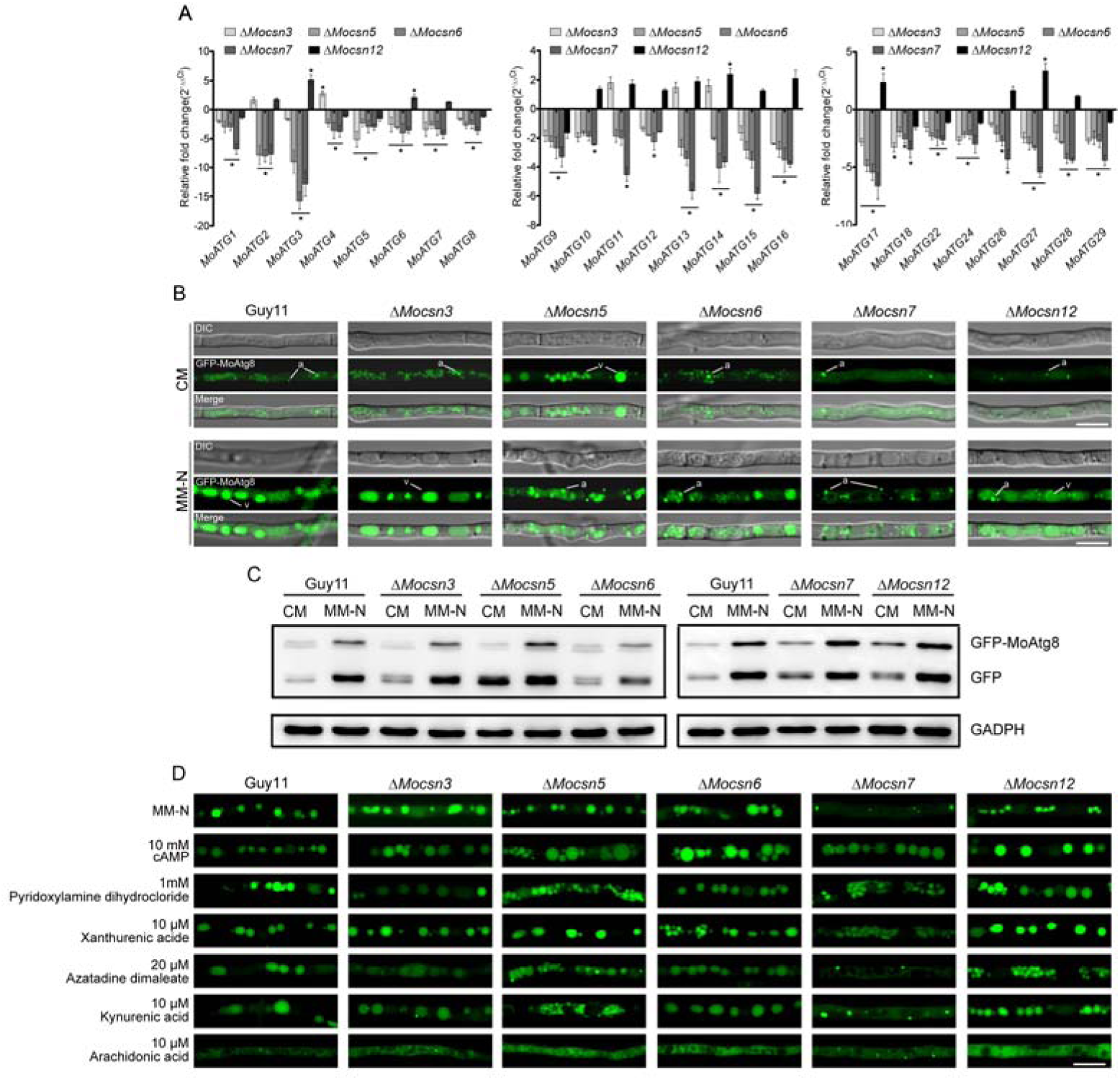
Targeted gene replacement of selected CSN subunits attenuated autophagic flux by cAMP biosynthesis and the expression of autophagy-associated proteins in the MoCSN defective strains. A, Relative fold-expression of genes coding autophagy-associated (ATG) proteins in the Δ*Mocsn3*, Δ*Mocsn5*, Δ*Mocsn6*, Δ*Mocsn7*, Δ*Mocsn12*, and wild-type strains after starvation for 6 h. B, Comparative localization pattern of the universal autophagy marker GFP-MoAtg8 fluorescence signals in the Δ*Mocsn3*, Δ*Mocsn5*, Δ*Mocsn6*, Δ*Mocsn7*, Δ*Mocsn12*, and wild-type strains under nutrient sufficient (CM) and nutrient deficient (MM-N) conditions. C, Immuno-blot-based analysis of GFP-MoAtg8/PE turn-over in the Δ*Mocsn3*, Δ*Mocsn5*, Δ*Mocsn6*, Δ*Mocsn7*, Δ*Mocsn12*, and wild-type strains under nutrient sufficient (CM) and nutrient deficient (MM-N) conditions. D, The micrograph showing the progression or suppression of autophagic flux in the Δ*Mocsn3*, Δ*Mocsn5*, Δ*Mocsn6*, Δ*Mocsn7*, Δ*Mocsn12*, and wild-type strains in MM-N supplemented independently with cAMP, pyridoxylamine, xanthurenic acid, azatadine dimaleate, kynurenic acid, and arachidonic acid. Note: the RT-qPCR assisted expression results were obtained from three biological replicates, each consisting of three technical replicates. The expression of the actin coding gene (MoACTIN) was used as the reference gene. Fold-expression of the MoATGs in the Δ*Mocsn* was computed using the delta delta-CT method (2^−ΔΔCT^). The expression of MoATGs in the wild-type strains was used as the internal control. For microscopy and immunoblotting assays, the individual strains were first cultured in CM for 2–3 days and later transferred into MM-N for 6 h to induce autophagy. The GADPH protein content was used as the control. (“a”) and (“v”) denote autophagosome and vacuole, respectively. Bar = 10 μm.

ATG8 proteins facilitate autophagosome formation, maturation, autophagosome-vacuolar fusion, and cargo selection by establishing anchorage with the headgroup of the membrane lipid phosphatidylethanolamine through lipidation(Martens and Fracchiolla, 2020). The association of ATG8 with all stages of the autophagic process makes it an ideal genetic marker for monitoring autophagic flux(Gómez-Sánchez et al., 2015). Therefore, to assess the initiation and progression of autophagic events in the Δ*Mocsn3*, Δ*Mocsn5*, Δ*Mocsn6*, Δ*Mocsn7*, and Δ*Mocsn12* strains relative to the wild-type, individual strains harboring the GFP-MoAtg8 fusion constructs were cultured under nutrient-sufficient (CM) and nutrient-deficient (MM-N) conditions and examined under a confocal microscope. Results obtained from microscopic examination showed the formation of autophagosomes (punctate structures of GFP-MoAtg8 fluorescence) in the Δ*Mocsn3*, Δ*Mocsn5*, Δ*Mocsn6*, and Δ*Mocsn12* strains under non-starvation conditions; there was no visible induction of autophagosome formation in the Δ*Mocsn7*, and the wild-type strains under nutrient-sufficient conditions (Figure 7B). Furthermore, we observed that targeted gene disruption of *MoCSN5, MoCSN6*, and *MoCSN12* substantially suppressed autophagic degradation under starvation, as puncta of GFP-MoAtg8 fluorescence remained visible in the autophago-vacuolar complex, whereas deletion of *MoCSN7* exclusively abolished autophagosome-vacuolar fusion under starvation (Figure 7B). Results obtained from western blot analyses conducted to measure autophagic flux in the individual strains further confirmed substantial suppression in the progression of autophagic degradation in the Δ*Mocsn5*, Δ*Mocsn7*, and Δ*Mocsn12* strains under starvation, whereas the levels of autophagic flux recorded in the Δ*Mocsn3* and Δ*Mocsn6* strains were comparable to the level of autophagic flux observed in the wild-type strain (Figure 7C). Meanwhile, of the metabolites that were downregulated in Δ*Mocsn3*, Δ*Mocsn5*, and Δ*Mocsn7*, exogenous application of cAMP exclusively restored autophagic flux-related defects observed in the Δ*Mocsn5*, Δ*Mocsn7*, and Δ*Mocsn12* strains (Figure 7D). Accordingly, we inferred that individual subunits of the CSN complex play diverse roles in the progression of autophagic flux in rice blast fungi. These observations suggest that the CSN complex contributes to the progression of autophagic flux by regulating cAMP generation and accumulation in *M. oryzae*.

## Discussion

The CSN signalosome complex was previously considered a conserved nuclear-localized protein complex(Füzesi-Levi et al., 2014). However, recent studies have shown that subunits of the CSN complex localize to both the nucleus and the cytoplasm in animal cell lines(Füzesi-Levi *et al*., 2014). All eight CSN subunits identified in rice blast fungus displayed a nucleocytoplasmic localization pattern. The uniform localization pattern of the CSN signalosome complex subunits in *M. oryzae* partly confirmed them as constituents of the CSN complex.

The presence of nuclear localization signals (NLS) or nuclear export signals (NES) facilitates the nucleocytoplasmic shuttling of proteins and protein complexes(Fu et al., 2018; Görlich et al., 1995; Johnson et al., 1999). The successful import or exit of proteins through the nuclear pore complex (NPC) is mediated by monopartite and bipartite proline-tyrosine-rich nuclear localization and leucine-rich nuclear export signals(Bernhofer et al., 2018). Karyopherin recognizes proteins containing these transport signal motifs, a conserved eukaryotic translocon that mediates the subsequent transport of nucleocytoplasmic-destined proteins into or out of the nucleus(Bernhofer *et al*., 2018).

Whether the CSN signalosome enters or exits the nucleus as a holocomplex; is still unknown. However, studies have shown that the NLS and NES in CSN5 facilitate its nucleocytoplasmic shuttling either as an individual subunit or as a mini-complex(Liu et al., 2010). In addition, these signals function in transporting cargo proteins into and out of the nucleus. For instance, studies have shown that truncating the NES signal in CSN5 impairs its role as an adaptor for cyclin-dependent kinase inhibitor 1 B (p27) and chromosomal maintenance 1/exportin 1 (CRM1), inhibits the nucleocytoplasmic translocation of p27-CSN5-CRM1, and prevents the degradation of tagged p27, which is linked to carcinogenic effects in humans(Sugiyama et al., 2001). However, the impact of the NLS/NES motif-containing subunits of the CSN complex and their likely impact on the nucleocytoplasmic localization of the CSN complex in filamentous fungi remains unknown.

In this study, we demonstrated that targeted replacement of translocon subunits, particularly *MoCSN5*, *MoCSN6*, and *MoCSN7*, suppressed growth and abolished asexual sporulation and pathogenicity of rice blast fungus. These observations provide practical insights into the significant contributions of the core translocon subunits of the CSN complex to the pathophysiological development of filamentous fungi.

In addition, we found that the NLS signal at the C-terminus of MoCsn7 functionally mediated the localization of MoCsn7 to the nucleus. Unexpectedly, we observed that fusing GFP to the C-terminus of MoCsn3 lacking the NLS failed to localize to the nucleoplasm. Given these observations, we speculate that MoCsn3 and MoCsn7, in association with nuclear translocons, likely regulate cytoplasm-nuclear migration of the CSN complex as a holocomplex or subcomplex.

The distinctive interaction pattern observed among subunits of the CSN complex from CoIP assays compared to the results obtained from Y2H assays suggests both direct and indirect interactions within subunits of the CSN complex in *M. oryzae.* Contrary to the suggestion that PCI domain-containing subunits play a conserved role in fostering interactions between subunits of the CSN complex(Lee et al., 2011; Qin et al., 2020), we demonstrated that targeted truncation of PCI in MoCsn1, MoCsn2, MoCsn3, MoCsn4, MoCsn7, and MoCsn12, and the MPN domain in MoCsn5 and MoCsn6 has no adverse or enhancing effect on the *in vitro* interaction network observed between subunits of the CSN complex in *M. oryzae.* Although MoCsn2 and MoCsn5 showed extensive one-on-one interaction with multiple subunits of the CSN complex according to results obtained from the Y2H bioassay conducted in the study, we postulated that MoCsn2, rather than MoCsn5, functions as a core regulator of the subunit-subunit interaction network in *M. oryzae.* Previous studies have shown that CSN5 actively interacts with the DNA-binding domain of GAL4 to produce a false-positive interaction outlook(Nordgård et al., 2001). However, the extent to which prevailing hierarchical *in vivo* and *in vitro* interaction networks recorded among CSN subunits influence the morphological and pathological development of rice blast is still unknown.

In addition to hierarchical interactions observed among individual subunits of the CSN complex, previous studies have demonstrated the existence of direct interactions between subunits of the COP9-signalosome complex and proteins outside the CSN complex (non-CSN complex proteins), including proteins associated with the ubiquitin-proteasome pathway, protein kinases, SOS1 protein, and eukaryotic translation initiation factors (eIFs)(Dubiel et al., 2015; Schwechheimer, 2004; Zarich et al., 2019). In humans, 825 proteins from diverse cellular pathways have been identified as putative interacting proteins in the COP9-signalosome complex(Fang et al., 2012). In this study, we recovered 575 proteins from the immunoprecipitation of the CSN subunits. However, only 57/575 core putative interactors (proteins present in the immuno-complexes of all the CSN subunits identified in *M. oryzae* except for MoCSN1) were identified. These include some known CNS complex-associated proteins, such as cullin1, cullin3, cullin 4 B, 26S proteasome non-ATPase regulatory subunit4, protein transporter SEC61 subunit alpha, E3 ubiquitin-protein ligase (UPL3), transcriptional repressor rco-1, E3-ubiquitin ligase complex SCF subunits con-3, 30S-ribosomal protein S14p/S29e, actin-like protein-3, AGC/AKT protein kinase, eukaryotic translation initiation factor2 subunit alpha, HET-C protein, and vacuolar protein sorting-associated protein-21. A total of 270 proteins were identified as common/core interactors of all CSN subunits identified in humans(Fang *et al*., 2012). The significant reduction in the number of core interactors identified for the CSN complex in *M. oryzae* compared to the CSN complex in humans likely reflects the differences in the complexity of cellular processes between filamentous fungi and animals.

Cyclooxygenases, lipoxygenases, and enzymes associated with the cytochrome P450 pathway mediate the generation of oxylipins, including the biosynthesis of prostaglandins, thromboxanes, and leukotrienes, through the peroxidation of polyunsaturated fatty acids (PUFA) such as arachidonic acid, linoleic acid, linolenic acid, eicosapentaenoic acid (EPA), stearidonic acid, adrenic acid, dihomo-γ-linolenic acid, and docosahexaenoic acid (DHA)(Gabbs et al., 2015; Tourdot et al., 2014). in In mammals, these biologically active PUFA derivatives (oxylipins) crucially regulate the progression of physiological, developmental, and cellular processes, including apoptosis, tissue repair, blood clotting, cell proliferation, blood vessel permeability, pain, inflammation, immune responses, and blood pressure(Buczynski et al., 2009). Irrespective of the significant importance of eicosanoids and their downstream derivatives in activating essential signaling pathways, under certain conditions, their presence in biological systems could have detrimental consequences(Gabbs *et al*., 2015).

Previous studies have shown that light period and temperature regulate membrane phospholipid and triacylglycerol metabolism. This synthesis increases the uptake of eicosanoids and their downstream derivatives(Norambuena et al., 2015). Recent studies have reported the generation of biologically active oxylipins derived from oxygenase (a significant type of cyclooxygenase) mediated oxidation of arachidonic acid in multiple human pathogenic fungal species, including *Candida albicans, Cryptococcus neoformans, Paracoccidioides brasiliensis, Fusarium dimerim, Microsporum audiouinii, Microsporum canis, Trichophyton rubrum, Absidia corymbifera, Histoplasma capsulatum, Blastomyces dermatitidis, Penicillium spp., Rhizopus spp., Rhizomucor pusillus*, and numerous *Aspergillus* species(Tourdot *et al*., 2014).

However, the photogenic role of the evolutionarily conserved constitutive photomorphogenesis complex in fungal physiology and pathogenesis and the possible regulation of PUFA and eicosanoid metabolism is still unknown. In this study, we experimentally demonstrated that targeted gene deletion of individual CSN signalosome subunits in *M. oryzae* abolished phototropism (photoresponse) in all defective strains except Δ*Mocsn3*. However, comparative non-targeted metabolomics analyses revealed a corresponding inhibition in the level (abundance) of two critical polyunsaturated fatty acids (eicosapentaenoic acid and arachidonic acid), phytosphingosine, prostaglandin b4, and accompanying derivative signaling molecules, including cAMP, indicating a potential breakdown in photo-dependent metabolism and signaling processes.

Oxylipins and oxylipin derivatives derived from oleic, linoleic, and linolenic acids function as essential chemical signaling molecules that modulate asexual and sexual reproduction in filamentous fungi, particularly in *A. nidulans*(*Tsitsigiannis et al., 2005*), there is no conclusive data regarding the direct or indirect regulatory influence of the CSN signalosome complex on the production of oxylipins and their derivatives in filamentous fungi(Tsitsigiannis and Keller, 2007). Targeted gene deletion of 3/5 of the NES/NLS motif-containing subunits (*MoCSN5*, *MoCSN6*, and *MoCSN7*) of the CSN complex in *M. oryzae* completely abolished asexual and sexual sporulation in rice blast fungus. These results directly correlate with the significant suppression in the metabolism of PUFAs, eicosanoids, and other oxylipin derivatives in the *MoCSN* defective strains generated in this study, especially the Δ*Mocsn7* strains.

Moreover, small changes in the biosynthesis of membrane lipids hinders the production of lipid-based signaling molecules required to stimulate the biosynthesis and uptake of amino acids. This suggests the existence of a close association between fatty acid metabolism and the cellular regulatory roles of the CSN complex. For instance, in plants, phenylalanine, tyrosine, and tryptophan synthesis pathways attenuate light-regulated redox homeostasis(Vivancos et al., 2011). Consistent with previous reports, we showed that in addition to the suppression of eicosanoid metabolism, targeted gene replacement of the CSN complex constituents in rice blast fungus inhibited the metabolism of diverse groups of amino acids, including glutamic acid, histidine, lysine, and arginine. We inferred from these results that the CSN signalosome complex likely plays a crucial role in fostering cross-talk between eicosanoid metabolism pathways, amino acid biosynthesis pathways, and nucleotide metabolism pathways.

Also, we demonstrated that the exogenous application of metabolites down-regulated in the metabolome of the defective strains, as well as metabolites identified exclusively in the metabolome of the Δ*Mocsn7* strain, partially restored conidiation defects recorded in Δ*Mocsn3*, Δ*Mocsn5*, Δ*Mocsn6*, and Δ*Mocsn12*, in a unanimous manner. The inability of these exogenously applied compounds to alter sporulation defects observed in the Δ*Mocsn7* strains, coupled with the hypersensitivity of the Δ*Mocsn7* strains to the cell wall and cell membrane stress-inducing osmolytes, support our hypothesis that Csn7 is likely the core subunit of the CSN signalosome complex functionally assigned to regulate the photo-responsive essential cellular metabolites needed to drive redox/oxidative balance and photosporogenesis in filamentous fungi. Therefore, the presence of the Csn7 subunit is essential for coordinating the activities of metabolic pathways in *M. oryzae* and other filamentous fungal species.

Small signaling molecules, including cAMP, kynurenic acid, xanthurenic acid, azatadine, and pyridoxyalamine, affect autophagic influx differently across organisms. For instance, studies have shown that cAMP functions as either a positive or negative regulator of autophagy, depending on the cell type(Grisan et al., 2021), kynurenic acid(Grisan *et al*., 2021) and pyridoxyalamine(Zhao et al., 2018) act as potent autophagic inhibitors, and xanthurenic acid has been shown to promote autophagosome biogenesis(Hou et al., 2020). Although the CSN has been implicated in the regulation of secondary metabolite biosynthesis and the progression of autophagic flux, there is no evidence that the CSN complex mediates the regulation of secondary metabolism during the progression of autophagic processes in filamentous fungi. Our findings showed that the targeted gene replacement *MoCSN7* exclusively suppressed the catabolism of arachidonic acid and consequently attenuated the generation of cAMP in the Δ*Mocsn3*, Δ*Mocsn5*, Δ*Mocsn6*, and Δ*Mocsn12* strains. In *S. cerevisiae*, elevated cAMP levels have been reported to suppress autophagic flux(Cebollero and Reggiori, 2009).

We also showed that exogenous application of cAMP fully restored autophagic flux in *MoCSN* defective strains. This observation is consistent with the positive role of cAMP in promoting autophagic flux in mammalian liver cells(Grisan *et al*., 2021; Wilson and Roach, 2002). Accordingly, we reasoned that the CSN signalosome complex, especially the Csn7 subunit, coordinates the initiation and progression of autophagic events in filamentous fungi under starvation conditions by facilitating an increase in the intracellular levels of cAMP, possibly via photo-dependent catabolism of arachidonic acid under nutrient-limited conditions.

## Conclusion

Although the COP9/CSN signalosome complex has long been identified as a positive regulator of autophagosome formation, maturation, and autophagic degradation, little is known about the molecular mechanisms that facilitate CSN-mediated regulation of autophagic flux, especially in filamentous fungi. Here, we observed that the genetic deactivation of putative translocon subunits of the CSN complex, particularly CSN7 (MoCsn7) in rice blast fungus, resulted in the accumulation of arachidonic acid exclusively in the *MoCSN7* mutant strains and caused an almost uniformly significant reduction in the generation of cAMP in the *MoCSN* null mutant strains generated in this study. Compared to the other *MoCSN* defective strains investigated, targeted replacement of *MoCSN7* exclusively abolished autophagic-vacuolar fusion and autophagic degradation under starvation. Phenocopy assays showed that the exogenous application of arachidonic acid inhibited autophagic flux in wild-type and defective strains under starvation conditions. In contrast, the exogenous application of cAMP fully rectified the abnormalities associated with the progression of autophagic flux in Δ*Mocsn* strains under starvation stress. Interestingly, the exogenous application of rapamycin, a macrolide antibiotic from *Streptomyces hygroscopicus*, is known to potently induce autophagy by suppressing the mTOR/TOR pathway in eukaryotes but failed to rescue autophagic defects observed in the Δ*Mocsn7* strains. While CSN5 has been suggested as a core regulator of the CSN complex in some selected organisms, here, we demonstrated that the CSN7a ortholog (*MoCSN7*) in rice blast fungus exerts a significant regulatory effect on the expression of autophagy-associated proteins and the progression of autophagic flux in *M. oryzae* through cAMP-dependent signal transduction. We showed that the CSN signalosome complex promotes morphological, reproductive, stress tolerance, and pathological development of rice blast fungus through synergistic and likely photo-dependent coordination of multiple cellular, biochemical, and metabolic pathways. Although cAMP has been shown to positively regulate pathogenic differentiation in fungi, the direct impact of cAMP on the progression of autophagic flux in filamentous fungi has not been reported. This study provides insights into the impact of CSN-mediated regulation of cAMP and other polyunsaturated fatty acids derivatives on autophagic degradation in *M. oryzae*. Understanding the regulatory parameters of pathogenesis in this pathogen is of great concern to the broader scientific community because it poses a huge economic burden.

## Materials and Methods

### Identification of CSN subunit *M. oryzae*, phylogenetic, domain structure, and sequence feature analyses

The eight genes coding for CSN subunits which comprises of the seven conical CSN subunits (Csn1-Csn7) and an additional subunit Csn12 were identified in the *M. oryzae* genome by performing a BLASTp/reverse BLASTp search with amino acid sequences retrieved for CSN complex subunits in *H. sapiens, N. crassa*, and *A. thaliana* from the KEGG and fungi and Oomycetes genomic resource platforms(Basenko et al., 2018). Domain profiling of the individual CSN subunits identified in the individual organisms using the Pfam 32.0 platform(Emms and Kelly, 2015). To identify MoCSN subunits with nuclear localization and Nuclear export signals (NLS/NES) and functionally evaluate or infer their influence on nucleocytoplasmic localization, we searched the amino acid sequences of the individual CSN subunits identified in *M. oryzae* using the NES prediction platform NetNES 1.1 Server(La Cour et al., 2004) and NLStradamus for NLS prediction(Nguyen Ba et al., 2009).

### Fungal strains and culture conditions

The parental wild-type *M. oryzae* (Guy11) strain, obtained from Dr. Didier Tharreau (CIRAD, Montpellier, France), was used as the background in the targeted gene replacement of all subunits of the *MoCSN* gene. Bacteria-competent cells used to propagate the constructed plasmids were prepared from *Escherichia coli* (*E. coli*) strain *DH5α*.

For vegetative growth, the wild-type, mutant, and complementation strains were cultured in a complete medium (for 1 L CM; 6 g yeast extract, 6 g casein hydrolysate, 10 g sucrose, and 20 g agar) at 25 ℃. For conidiation, the strains were cultured on rice bran agar medium (for 1 L RBA; 40 g rice bran, 20 g agar, pH 6.0) for 7 days under dark conditions. The cultured strains were later transferred into an incubator with continuous light for 3 days before vegetative hyphae were removed using sterilized microslides. Stress sensitivity of the individual strains was assayed by culturing the mutant strains along with the wild-type strains on CM supplemented with different stress-inducing agents (oxidative stress-inducing agents: 200 μg/mL Calcofluor White (F3543; Sigma-Aldrich, St. Louis, MO, USA), 0.7 M NaCl, 0.01% sodium dodecyl sulfate, 200 μg/mL Congo Red (0379; TAGENE, Xiamen, China); reductive stress-inducing agents: 2 mM DTT (1758-9030; Inalco, Paris, France).

For the circadian rhythm assay, the wild-type and Δ*Mocsn* strains were cultured on prune agar (PA) medium and incubated for 2 days under dark conditions at a stable temperature of 25 ℃. After 2 days, the strains were transferred into an incubator with a photoperiod of 12 h light/12 h dark. Note: PA media with a final pH of 6.5 was prepared using 40 mL prune juice, 2.5 g lactose, 2.5 g sucrose, 1 g yeast extract, and 20 g agar.

### Generation of gene replacement mutant and complementation

Split markers for gene knockout were constructed and used for targeted gene replacement of *MoCSN* in *M. oryzae*. To construct split markers for *MoCSN3*, 0.97 kb upstream and 1.29 kb downstream fragments of the flanking regions were amplified with primers *MoCSN3*-AF/AR and *MoCSN3*-BF/BR, respectively; for *MoCSN5*, 1.07 kb upstream and 1.26 kb downstream fragments of the flanking regions were amplified with primers *MoCSN5*-AF/AR and *MoCSN5*-BF/BR, respectively; for *MoCSN6*, 1.41 kb upstream and 0.97 kb downstream fragments of the flanking regions were amplified with primers *MoCSN6*-AF/AR and *MoCSN6*-BF/BR, respectively; *MoCSN7*, 1.12 kb upstream and 0.74 kb downstream fragments of the flanking regions were amplified with primers *MoCSN7*-AF/AR and *MoCSN7*-BF/BR, respectively; *MoCSN12*, 0.87 kb upstream and 0.98 kb downstream fragments of the flanking regions were amplified with primers *MoCSN12*-AF/AR and *MoCSN12*-BF/BR, respectively.

The upstream fragments were cloned into the upstream half of HPH on pCX62 using the *Kpn* I and *Eco*R I restriction enzyme sites. The downstream fragments were cloned into the downstream half of HPH on the pCX62 vector using the *Bam*H I and *Xba* I restriction enzyme sites and overlap extension PCR cloning (OE-PCR)(Bryksin and Matsumura, 2013). The products used to construct the targeted gene deletion constructs were amplified using primer pairs MoCSN-AF+HY/R and YG/F+ MoCSN-BR listed in (Table S5). *M. oryzae* protoplast preparation and fungal transformation were performed as described by(Nakayashiki et al., 2005; Talbot et al., 1993). The transformants were screened using MoCSN-OF/MoCSN-OR and MoCSN-UF/MoCSN-UR. The MoCSN gene-defective strains generated in this study were confirmed using a Southern blotting assay.

To construct the complementation/MoCSN-GFP-fusion vector, the fragment, including the native promoter and the whole ORF sequence without a stop codon, was amplified. The product was cloned into the pKNTG vector upstream of the GFP site using the *Eco*R I and *Bam*H I enzyme restriction sites. To generate complementation and localization strains, the constructed vectors for the individual genes were transformed into protoplasts of the respective Δ*Mocsn* strains. The transformants were screened using PCR and the MoCSN-OF and GFP-R primer pair and scanned for fluorescence intensity using a microscope.

### Genomic DNA isolation

Genomic DNA extraction from wild-type Guy11, Δ*Mocsn*, and complementation strains using the CTAB DNA extraction procedure described by (Brandfass and Karlovsky, 2008; Shabbir et al., 2022) with minor modifications.

### Conidiation, appressoria formation, conidiophore assessment, and mating-type assays

Conidiation was evaluated by washing conidia from 10-day-old culture plates with sterilized ddH_2_O and filtered through three layers of lens paper into a 2 mL EP tube. Appressorium formation was monitored by placing 20 μL of spore suspension (concentration of 5×10^4^ spores per mL) on hydrophobic coverslips and incubating under humid and dark conditions and at a temperature of 28 ℃ for 4, 8, 16, and 24 h. An average of 100 conidia was examined for each experiment, and consistent results from three independent biological experiments with three technical replicates were used for appressorium formation computation.

For the conidiophore assay, blocks of RBA containing the strains were transferred onto microslides, with the side bearing the hyphae resting on the slide surface, and incubated at 28℃ in the light for 48 h and 96 h. After this period, the blocks were removed and treated with lactophenol cotton blue (LCB) dye for 5 min. A 100 mL LCB solution (20 mL phenol, 0.6 g cotton blue, 44 mL glycerin, 16 mL lactic acid, and ddH_2_O) was diluted three-fold to obtain a working solution. Finally, the excess dye was washed off the slides with ddH_2_O and visualized using the bright field (BF) mode of an Olympus Bx51 Microscope.

Mating capabilities of the individual strains were assayed by pairing or co-culturing the Δ*Mocsn* with the standard tester strain KA3 (*MAT1-1)* cultivated on oatmeal agar medium (OA) at a stable temperature of 20℃ for 3-4 weeks. The KA3 (*MAT1-1)* standard tester strains were obtained from Dr. Didier Tharreau.

### Infection assay

For hyphae-mediated infection assessment, the individual strains were cultured in liquid CM for 3 days in a shaking incubator at a rotator speed of 110 rpm and a stable temperature of 28℃. The mycelia were filtered using Whatman filter paper, washed with sterilized ddH_2_O, and allowed to stand for 5 min until excess water was drained. The mycelia were then used as propagules to inoculate intact and injured leaves excised from 7-day-old barley seedlings. The inoculated leaves and the uninoculated controls were first incubated in a dark chamber with 90% relative humidity and a temperature of 25℃ for 24 h. The inoculated leaves along with control group were transferred to a growth chamber with a photoperiod of 12 h light/12 h dark. The mycelia obtained from the individual strains were also ground to prepare mycelial suspensions. The mycelial suspensions were fortified with 0.02% v/v Tween 20. Mycelial suspensions were used to spray-inoculate 3-week-old blast-susceptible rice seedlings (*Oryza sativa* cv. CO39), following the procedures and inoculation conditions described for barley infection. Disease development and lesion severity were assessed at 7 days post-inoculation (dpi) and used as a measure of pathogenicity and virulence characteristics of the defective strains compared to the wild-type.

Histopathological examinations (host penetration and colonization assays) were performed by inoculating the underside of the barley leaves with mycelia. Inoculated tissues were incubated under the conditions described above. Host invasion and colonization characteristics of the individual strains were assessed at 24 hours post-inoculation (hpi) using a light microscope.

### Microscopy analysis

For microscopy, an Olympus DP72 fluorescent microscope or a Nikon A1 plus confocal microscope was used to observe the fluorescence of GFP and mCherry. The emission and excitation wavelengths were 488 nm and 561 nm, respectively.

### Co-localization assay

To confirm the localization of MoCsn, we constructed a MoCsn-GFP vector and co-transformed MoCsn-GFP with the histone-mCherry (His-mCherry) marker into Guy11. The transformants were screened using PCR and a primer pair (MoCSN-OF and GFP-R) and further confirmed using fluorescence microscopy. Histone-mCherry (His-mCherry)(Zhang et al., 2019) was obtained from Dr. Lianhu Zhang of Jiangxi Agricultural University.

### Transmission electron microscopy observation

Vegetative hyphae of the Guy11 and Δ*Mocsn* strains were cultured in liquid CM for 3 days. The hyphae were fixed with 2.5% glutaraldehyde in phosphate buffer (pH 7.0), washed three times with phosphate buffer, fixed with 1% OsO_4_ in phosphate buffer for 1 h, and washed three times with phosphate buffer. The specimens were dehydrated using a graded series of ethanol (30%, 50%, 80%, 90%, 95%, and 100%) for approximately 20 min at each concentration, following previously reported procedures(Zhong et al., 2016).

### Co-Immunoprecipitation assay

Total proteins extracted from the individual MoCsn-GFP strains, along with the control strain harboring the empty GFP vector, were incubated with 30 μL of anti-GFP magarose beads (beads No.SM03801; Smart-life Sciences, China) for 4 h at 4 ℃ to immunoprecipitate the GFP-fusion proteins from cellular extracts. Then, a magnetic frame was used to wash the beads three times with 500 μL cold wash buffer (50 mM Tris, 0.15 M NaCl, pH 7.4) and resuspended in 80 μL SDS-loading buffer. Proteins eluted from anti-GFP magarose beads were analyzed using immunoblotting with anti-GFP antibodies (anti-GFP No.334578; Abmart, China), followed by mass spectrometry (BGI, China).

### Yeast two-hybrid assay

Full-length MoCsn cDNA was amplified and cloned into a pGBKT7/pGADT7 plasmid to obtain the bait vector MoCsn-BD and prey vector MoCsn-AD according to a previous protocol(Young, 1998) to generate clone vectors and positive transgenic yeast strains for yeast two-hybrid screening of interactions between the subunits of the MoCsn complex. The interaction between pGBKT7-53 and pGADT7-T was used as a positive control, and pGBKT7-Lam and pGADT7-T were used as negative control. The resultant bait and prey vectors were confirmed using sequencing and co-transformation into the yeast strain AH109. All transformants were assayed with 1×10^6^ cells/μL droplets on SD-Leu-Trp and SD-Leu-Trp-His-Ade plates with 20 mg/mL X-α-gal.

### Total RNA extraction and Real-time RT-PCR assay

The strains were grown in liquid CM in a shaking incubator at a stable rotor speed of 110 rpm for 4 days. Mycelia from the cultured strains were filtered and washed thoroughly with sterilized ddH_2_O. Excess water was removed using Whatman filter paper, and the mycelia were freeze-dried in liquid nitrogen. The samples were ground in liquid nitrogen, and total RNA was extracted using the HiPure Universal RNA kit (R4130-02; Magen, China). The expression of individual genes under different conditions or in Δ*Mocsn* was monitored using quantitative real-time PCR (qRT-PCR) assays. Reverse RNA transcription was performed using the PrimeScript RT Reagent Kit with gDNA Eraser (RR047A; Takara Bio, Japan). A 10 μL reaction mix was formulated as follows: 5 μL TB green, 3.4 μL RNase-free water, 0.3 μL of each 10 μM forward and reverse primers (listed in Table S5), and 1 μL cDNA template. qRT-PCR was performed using an Eppendorf Realplex2 Master Cycler (Eppendorf AG 223341, Hamburg, Germany). Raw qRT-PCR data were analyzed using the delta delta-CT (2^-ΔΔCt^) method described by(Rao et al., 2013; Sami et al., 2020). Tubulin expression was used as the reference or internal control. Error bars represent the mean ± SD. RT-qPCR data were generated from three independent biological experiments, each consisting of three technical replicates.

### Metabolomics analysis

The strains were cultured in liquid CM for 4 days in a shaking incubator at a stable rotor speed of 110 rpm. The mycelia were filtered, quickly frozen in liquid nitrogen, and then kept in a low-temperature freeze dryer (Labconco Free Zone 12 L, Labconco, USA) to dry for 48 h. The dry powder (0.025 g) was placed in a 1.5 mL Eppendorf tube containing 800 μL extraction buffer (methanol: acetonitrile: ddH2O, 2:2:1 (v/v/v)), 50 Hz for 5 min, and incubated in an ultrasonic water bath for 10 min.

After incubation, the mixture was centrifuged at 15294 g for 15 min. The supernatants were then pipetted into new 1.5 mL sterilized Eppendorf tubes and concentrated using vacuum-assisted freezing until dry. Then, 200 μL of 10% methanol was added and incubated in an ultrasonic water bath for 10 min to dissolve the precipitate. After incubation, the mixture was centrifuged at 15294 g for 15 min, filtered through 0.22 μm Milex Millipore membranes into sample bottles with glass inserts, and stored at 4 ℃. Non-targeted metabolomics analysis by BGI, China, was used in this study.

## Acknowledgements

We are grateful to the members of Z.W. laboratory for their insightful discussions.

## AUTHOR CONTRIBUTIONS

J.N. and L.L*, conceived, designed and sourced funding for the research, J.N., L.L*, and Z.W, secured the funding, L.L, J.N., H.G., W.B., J.C., S.R.A., and Q.A. performed the experiments. L.L*., W.Z. J.N., and H.G. analyzed the data. L.L., drafted the manuscript. J.N., L.L, Q.C., and Z.W. revised the manuscript. All authors contributed to the final manuscript.

## Availability of data and materials

The data supporting the findings of this study are available from the corresponding author upon reasonable request.

## Declarations

Ethics Approval and Consent to Participate in this study complied with the ethical standards of China where this research was carried out.

Consent for Publication: All authors consent to publication of the study. Competing interests: The authors declare that they have no competing interests.

## Supplementary Information

**Figure S1:** The carboxyl-terminal (C-terminal) is essential for nucleocytoplasmic localization of MoCsn3 and MoCsn7 during vegetative growth and pathogenic differentiation in *M. oryzae*.

**Figure S2:** Southern blotting assays confirmed the replacement of genes coding for NLS/NES subunits of the CSN complex in *M. oryzae* via single insertion and successful complementation.

**Figure S3:** Re-introduction of full-length ORF of individual MoCSN genes rescued growth and conidiation defects associated with Δ*Mocsn strains*.

**Figure S4:** Targeted gene disruption of *MoCSN7* severely compromised cell wall integrity of *M. oryzae*.

**Figure S5:** The expression pattern of *MoCSN*, ubiquitinating, and phytochrome-related genes in the *M. oryzae* strains treated with blue and red light.

**Figure S6:** The PCA analysis of the reliability of metabolites identified between replicates for the individual strains.

**Figure S7:** The enrichment analysis of metabolic pathways of differential metabolites in Guy11, Δ*Mocsn3*, Δ*Mocsn5*, and Δ*Mocsn7*.

**Table S1:** Annotation and Q-score values for MoCsn interaction subunits recovered from the individual MoCsn immuno-complexes.

**Table S2:** The numerical distribution of metabolites identified in the metabolome of individual strains.

**Table S3:** Exogenous application down-regulated metabolites selectively rescued vegetative growth and conidiation defects Δ*Mocsn3*, Δ*Mocsn5*, Δ*Mocsn6*, and Δ*Mocsn12* strains.

**Table S4:** List and gene accession numbers of autophagy-related proteins identified *M. oryzae*.

**Table S5:** List and nucleotide sequence for the primer pairs used in this study.

## References

Barth, E., Hübler, R., Baniahmad, A., and Marz, M. (2016). The evolution of COP9 signalosome in unicellular and multicellular organisms. Genome Biology and Evolution 8, 1279–1289.

Basenko, E.Y., Pulman, J.A., Shanmugasundram, A., Harb, O.S., Crouch, K., Starns, D., Warrenfeltz, S., Aurrecoechea, C., Stoeckert Jr, C.J., and Kissinger, J.C. (2018). FungiDB: an integrated bioinformatic resource for fungi and oomycetes. Journal of Fungi 4, 39.

Bernhofer, M., Goldberg, T., Wolf, S., Ahmed, M., Zaugg, J., Boden, M., and Rost, B. (2018). NLSdb—major update for database of nuclear localization signals and nuclear export signals. Nucleic acids research 46, D503–D508.

Brandfass, C., and Karlovsky, P. (2008). Upscaled CTAB-based DNA extraction and real-time PCR assays for Fusarium culmorum and F. graminearum DNA in plant material with reduced sampling error. International journal of molecular sciences 9, 2306–2321.

Bryksin, A., and Matsumura, I. (2013). Overlap extension PCR cloning. Synthetic Biology, 31–42.

Buczynski, M., Dumlao, D., and Dennis, E. (2009). An integrated omics analysis of eicosanoid biology (vol 50, pg 1015, 2009). Journal of lipid research 50, 1505–1505.

Busch, S., Eckert, S.E., Krappmann, S., and Braus, G.H. (2003). The COP9 signalosome is an essential regulator of development in the filamentous fungus Aspergillus nidulans. Molecular microbiology 49, 717–730.

Cebollero, E., and Reggiori, F. (2009). Regulation of autophagy in yeast Saccharomyces cerevisiae. Biochimica et Biophysica Acta (BBA)-Molecular Cell Research 1793, 1413–1421.

Chamovitz, D., and Deng, X.W. (1997). The COP9 complex: a link between photomorphogenesis and general developmental regulation? Plant, Cell & Environment 20, 734–739.

Chen, Y., Shao, X., Cao, J., Zhu, H., Yang, B., He, Q., and Ying, M. (2021). Phosphorylation regulates cullin-based ubiquitination in tumorigenesis. Acta Pharmaceutica Sinica B 11, 309–321.

Cope, G.A., and Deshaies, R.J. (2003). COP9 signalosome: a multifunctional regulator of SCF and other cullin-based ubiquitin ligases. Cell 114, 663–671.

Dong, B., Liu, X.-H., Lu, J.-P., Zhang, F.-S., Gao, H.-M., Wang, H.-K., and Lin, F.-C. (2009). MgAtg9 trafficking in Magnaporthe oryzae. Autophagy 5, 946–953.

Dubiel, D., Rockel, B., Naumann, M., and Dubiel, W. (2015). Diversity of COP9 signalosome structures and functional consequences. FEBS letters 589, 2507–2513.

Emms, D.M., and Kelly, S. (2015). OrthoFinder: solving fundamental biases in whole genome comparisons dramatically improves orthogroup inference accuracy. Genome biology 16, 1–14.

Fang, L., Kaake, R.M., Patel, V.R., Yang, Y., Baldi, P., and Huang, L. (2012). Mapping the protein interaction network of the human COP9 signalosome complex using a label-free QTAX strategy. Molecular & Cellular Proteomics 11, 138–147.

Fu, X., Liang, C., Li, F., Wang, L., Wu, X., Lu, A., Xiao, G., and Zhang, G. (2018). The rules and functions of nucleocytoplasmic shuttling proteins. International journal of molecular sciences 19, 1445.

Füzesi-Levi, M.G., Ben-Nissan, G., Bianchi, E., Zhou, H., Deery, M.J., Lilley, K.S., Levin, Y., and Sharon, M. (2014). Dynamic regulation of the COP9 signalosome in response to DNA damage. Molecular and cellular biology 34, 1066–1076.

Füzesi-Levi, M.G., Fainer, I., Ivanov Enchev, R., Ben-Nissan, G., Levin, Y., Kupervaser, M., Friedlander, G., Salame, T.M., Nevo, R., and Peter, M. (2020). CSNAP, the smallest CSN subunit, modulates proteostasis through cullin-RING ubiquitin ligases. Cell Death & Differentiation 27, 984–998.

Gabbs, M., Leng, S., Devassy, J.G., Monirujjaman, M., and Aukema, H.M. (2015). Advances in our understanding of oxylipins derived from dietary PUFAs. Advances in nutrition 6, 513–540.

Gómez-Sánchez, R., Pizarro-Estrella, E., Yakhine-Diop, S.M., Rodríguez-Arribas, M., Bravo-San Pedro, J.M., Fuentes, J.M., and González-Polo, R.A. (2015). Routine Western blot to check autophagic flux: cautions and recommendations. Analytical biochemistry 477, 13–20.

Görlich, D., Kostka, S., Kraft, R., Dingwall, C., Laskey, R.A., Hartmann, E., and Prehn, S. (1995). Two different subunits of importin cooperate to recognize nuclear localization signals and bind them to the nuclear envelope. Current Biology 5, 383–392.

Grisan, F., Iannucci, L.F., Surdo, N.C., Gerbino, A., Zanin, S., Di Benedetto, G., Pozzan, T., and Lefkimmiatis, K. (2021). PKA compartmentalization links cAMP signaling and autophagy. Cell Death & Differentiation 28, 2436–2449.

Gutierrez, C., Chemmama, I.E., Mao, H., Yu, C., Echeverria, I., Block, S.A., Rychnovsky, S.D., Zheng, N., Sali, A., and Huang, L. (2020). Structural dynamics of the human COP9 signalosome revealed by cross-linking mass spectrometry and integrative modeling. Proceedings of the National Academy of Sciences 117, 4088–4098.

Hartman, J.J., Mahr, J., McNally, K., Okawa, K., Iwamatsu, A., Thomas, S., Cheesman, S., Heuser, J., Vale, R.D., and McNally, F.J. (1998). Katanin, a microtubule-severing protein, is a novel AAA ATPase that targets to the centrosome using a WD40-containing subunit. Cell 93, 277–287.

He, Q., Cheng, P., He, Q., and Liu, Y. (2005). The COP9 signalosome regulates the Neurospora circadian clock by controlling the stability of the SCFFWD-1 complex. Genes & development 19, 1518–1531.

Hou, T., Sun, X., Zhu, J., Hon, K.-L., Jiang, P., Chu, I.M.-T., Tsang, M.S.-M., Lam, C.W.-K., Zeng, H., and Wong, C.-K. (2020). IL-37 ameliorating allergic inflammation in atopic dermatitis through regulating microbiota and AMPK-mTOR signaling pathway-modulated autophagy mechanism. Frontiers in Immunology 11, 752.

Johnson, C., Van Antwerp, D., and Hope, T.J. (1999). An N-terminal nuclear export signal is required for the nucleocytoplasmic shuttling of IκBα. The EMBO journal 18, 6682–6693.

Kim, W.D., Mathavarajah, S., and Huber, R.J. (2022). The cellular and developmental roles of cullins, neddylation, and the COP9 signalosome in Dictyostelium discoideum. Frontiers in Physiology, 311.

Klionsky, D.J., and Eskelinen, E.-L. (2014). The vacuole vs. the lysosome: When size matters. Autophagy 10, 185–187.

La Cour, T., Kiemer, L., Mølgaard, A., Gupta, R., Skriver, K., and Brunak, S. (2004). Analysis and prediction of leucine-rich nuclear export signals. Protein Engineering Design and Selection 17, 527–536.

Lee, M.-H., Zhao, R., Phan, L., and Yeung, S.-C.J. (2011). Roles of COP9 signalosome in cancer. Cell cycle 10, 3057–3066.

Lee, Y. (2020). Regulation and function of capicua in mammals. Experimental & molecular medicine 52, 531–537.

Licursi, V., Salvi, C., De Cesare, V., Rinaldi, T., Mattei, B., Fabbri, C., Serino, G., Bramasole, L., Zimbler, J.Z., and Pick, E. (2014). The COP9 signalosome is involved in the regulation of lipid metabolism and of transition metals uptake in Saccharomyces cerevisiae. The FEBS journal 281, 175–190.

Liu, Q., Yu, J., Zhuo, X., Jiang, Q., and Zhang, C. (2010). Pericentrin contains five NESs and an NLS essential for its nucleocytoplasmic trafficking during the cell cycle. Cell research 20, 948–962.

Martens, S., and Fracchiolla, D. (2020). Activation and targeting of ATG8 protein lipidation. Cell discovery 6, 23.

Moghe, S., Jiang, F., Miura, Y., Cerny, R.L., Tsai, M.Y., and Furukawa, M. (2011). The CUL3-KLHL18 ligase regulates mitotic entry and ubiquitylates Aurora-A. Biology open 1, 82–91.

Nakayashiki, H., Hanada, S., Quoc, N.B., Kadotani, N., Tosa, Y., and Mayama, S. (2005). RNA silencing as a tool for exploring gene function in ascomycete fungi. Fungal Genetics and Biology 42, 275–283.

Nguyen Ba, A.N., Pogoutse, A., Provart, N., and Moses, A.M. (2009). NLStradamus: a simple Hidden Markov Model for nuclear localization signal prediction. BMC bioinformatics 10, 1–11.

Norambuena, F., Morais, S., Emery, J.A., and Turchini, G.M. (2015). Arachidonic acid and eicosapentaenoic acid metabolism in juvenile Atlantic salmon as affected by water temperature. PLoS One 10, e0143622.

Nordgård, O., Dahle, Ø., Andersen, T.Ø., and Gabrielsen, O.S. (2001). JAB1/CSN5 interacts with the GAL4 DNA binding domain: a note of caution about two-hybrid interactions. Biochimie 83, 969–971.

Norvienyeku, J., Lin, L., Waheed, A., Chen, X., Bao, J., Aliyu, S.R., Lin, L., Shabbir, A., Batool, W., and Zhong, Z. (2021). Bayogenin 3-O-cellobioside confers non-cultivar-specific defence against the rice blast fungus Pyricularia oryzae. Plant Biotechnology Journal 19, 589–601.

Oberhauser, L., Granziera, S., Colom, A., Goujon, A., Lavallard, V., Matile, S., Roux, A., Brun, T., and Maechler, P. (2020). Palmitate and oleate modify membrane fluidity and kinase activities of INS-1E β-cells alongside altered metabolism-secretion coupling. Biochimica et Biophysica Acta (BBA)-Molecular Cell Research 1867, 118619.

Osterlund, M.T., Ang, L.-H., and Deng, X.W. (1999). The role of COP1 in repression of Arabidopsis photomorphogenic development. Trends in cell biology 9, 113–118.

Peng, Z., Serino, G., and Deng, X.-W. (2001). A role of Arabidopsis COP9 signalosome in multifaceted developmental processes revealed by the characterization of its subunit 3.

Pintard, L., Kurz, T., Glaser, S., Willis, J.H., Peter, M., and Bowerman, B. (2003). Neddylation and deneddylation of CUL-3 is required to target MEI-1/Katanin for degradation at the meiosis-to-mitosis transition in C. elegans. Current Biology 13, 911–921.

Qin, N., Xu, D., Li, J., and Deng, X.W. (2020). COP9 signalosome: Discovery, conservation, activity, and function. Journal of Integrative Plant Biology 62, 90–103.

Rao, X., Huang, X., Zhou, Z., and Lin, X. (2013). An improvement of the 2^ (–delta delta CT) method for quantitative real-time polymerase chain reaction data analysis. Biostatistics, bioinformatics and biomathematics 3, 71.

Sami, A., Naqvi, S.S., Qayyum, M., Rao, A.R., Sabitaliyevich, U.Y., and Ahmad, M.S. (2020). Calcium based siRNA coating: a novel approach for knockdown of HER2 gene in MCF-7 cells using gold nanoparticles. Cellular and Molecular Biology 66, 105–111.

Schwechheimer, C. (2004). The COP9 signalosome (CSN): an evolutionary conserved proteolysis regulator in eukaryotic development. Biochimica et Biophysica Acta (BBA)-Molecular Cell Research 1695, 45–54.

Serino, G., and Deng, X.-W. (2003). The COP9 signalosome: regulating plant development through the control of proteolysis. Annual review of plant biology 54, 165–182.

Shabbir, A., Batool, W., Yu, D., Lin, L., An, Q., Xiaomin, C., Guo, H., Yuan, S., Malota, S., and Wang, Z. (2022). Magnaporthe oryzae Chloroplast Targeting Endo-β-1, 4-Xylanase I MoXYL1A Regulates Conidiation, Appressorium Maturation and Virulence of the Rice Blast Fungus. Rice 15, 44.

Shackleford, T.J., and Claret, F.X. (2010). JAB1/CSN5: a new player in cell cycle control and cancer. Cell division 5, 1–14.

Smith, P., Leung-Chiu, W., Montgomery, R., Orsborn, A., Kuznicki, K., Gressman-Coberly, E., Mutapcic, L., and Bennett, K. (2002). The GLH proteins, Caenorhabditis elegans P granule components, associate with CSN-5 and KGB-1, proteins necessary for fertility, and with ZYX-1, a predicted cytoskeletal protein. Developmental Biology 251, 333–347.

Sugiyama, Y., Tomoda, K., Tanaka, T., Arata, Y., Yoneda-Kato, N., and Kato, J.-y. (2001). Direct binding of the signal-transducing adaptor Grb2 facilitates down-regulation of the cyclin-dependent kinase inhibitor p27Kip1. Journal of Biological Chemistry 276, 12084–12090.

Suisse, A., Békés, M., Huang, T.T., and Treisman, J.E. (2018). The COP9 signalosome inhibits Cullin-RING E3 ubiquitin ligases independently of its deneddylase activity. Fly 12, 118–126.

Talbot, N.J., Ebbole, D.J., and Hamer, J.E. (1993). Identification and characterization of MPG1, a gene involved in pathogenicity from the rice blast fungus Magnaporthe grisea. The Plant Cell 5, 1575–1590.

Tourdot, B.E., Ahmed, I., and Holinstat, M. (2014). The emerging role of oxylipins in thrombosis and diabetes. Frontiers in pharmacology 4, 176.

Tsitsigiannis, D.I., and Keller, N.P. (2007). Oxylipins as developmental and host– fungal communication signals. Trends in microbiology 15, 109–118.

Tsitsigiannis, D.I., Kowieski, T.M., Zarnowski, R., and Keller, N.P. (2005). Three putative oxylipin biosynthetic genes integrate sexual and asexual development in Aspergillus nidulans. Microbiology 151, 1809–1821.

Vivancos, P.D., Driscoll, S.P., Bulman, C.A., Ying, L., Emami, K., Treumann, A., Mauve, C., Noctor, G., and Foyer, C.H. (2011). Perturbations of amino acid metabolism associated with glyphosate-dependent inhibition of shikimic acid metabolism affect cellular redox homeostasis and alter the abundance of proteins involved in photosynthesis and photorespiration. Plant physiology 157, 256–268.

Wang, M., Yang, X., Ruan, R., Fu, H., and Li, H. (2018). Csn5 is required for the conidiogenesis and pathogenesis of the Alternaria alternata tangerine pathotype. Frontiers in Microbiology 9, 508.

Wang, X., Li, W., Piqueras, R., Cao, K., Deng, X.W., and Wei, N. (2009). Regulation of COP1 nuclear localization by the COP9 signalosome via direct interaction with CSN1. The Plant Journal 58, 655–667.

Wilson, A.M., Wilken, P.M., Wingfield, M.J., and Wingfield, B.D. (2021). Genetic networks that govern sexual reproduction in the Pezizomycotina. Microbiology and Molecular Biology Reviews 85, e00020–00021.

Wilson, W.A., and Roach, P.J. (2002). Nutrient-regulated protein kinases in budding yeast. Cell 111, 155–158.

Young, K. (1998). Yeast two-hybrid: so many interactions,(in) so little time….Biology of reproduction 58, 302–311.

Yu, Z., Kleifeld, O., Lande-Atir, A., Bsoul, M., Kleiman, M., Krutauz, D., Book, A., Vierstra, R.D., Hofmann, K., and Reis, N. (2011). Dual function of Rpn5 in two PCI complexes, the 26S proteasome and COP9 signalosome. Molecular biology of the cell 22, 911–920.

Zarich, N., Anta, B., Fernández-Medarde, A., Ballester, A., de Lucas, M.P., Cámara, A.B., Anta, B., Oliva, J.L., Rojas-Cabañeros, J.M., and Santos, E. (2019). The CSN3 subunit of the COP9 signalosome interacts with the HD region of Sos1 regulating stability of this GEF protein. Oncogenesis 8, 2.

Zhang, L., Zhang, D., Chen, Y., Ye, W., Lin, Q., Lu, G., Ebbole, D.J., Olsson, S., and Wang, Z. (2019). Magnaporthe oryzae CK2 accumulates in nuclei, nucleoli, at septal pores and forms a large ring structure in appressoria, and is involved in rice blast pathogenesis. Frontiers in cellular and infection microbiology 9, 113.

Zhao, X., Chen, Y., Tan, X., Zhang, L., Zhang, H., Li, Z., Liu, S., Li, R., Lin, T., and Liao, R. (2018). Advanced glycation end-products suppress autophagic flux in podocytes by activating mammalian target of rapamycin and inhibiting nuclear translocation of transcription factor EB. The Journal of Pathology 245, 235–248.

Zhong, K., Li, X., Le, X., Kong, X., Zhang, H., Zheng, X., Wang, P., and Zhang, Z. (2016). MoDnm1 dynamin mediating peroxisomal and mitochondrial fission in complex with MoFis1 and MoMdv1 is important for development of functional appressorium in Magnaporthe oryzae. PLoS pathogens 12, e1005823.

Zhou, Z., Wang, Y., Cai, G., and He, Q. (2012). Neurospora COP9 signalosome integrity plays major roles for hyphal growth, conidial development, and circadian function. PLoS Genetics 8, e1002712.

